# Spatial Relationships Matter: Kinesin-1 Molecular Motors Transport Liposome Cargo Through 3D Microtubule Intersections *In Vitro*

**DOI:** 10.1101/2023.12.01.569616

**Authors:** Brandon M Bensel, Samantha Previs, Carol Bookwalter, Kathleen M Trybus, Sam Walcott, David M Warshaw

## Abstract

Kinesin-1 ensembles maneuver vesicular cargoes through intersections in the 3-dimensional (3D) intracellular microtubule (MT) network. To characterize directional outcomes (straight, turn, terminate) at MT intersections, we challenge 350 nm fluid-like liposomes transported by ∼10 constitutively active, truncated kinesin-1 KIF5B (K543) with perpendicular 2-dimensional (2D) and 3D intersections *in vitro*. Liposomes frequently pause at 2D and 3D intersections (∼2s), suggesting that motor teams can simultaneously engage each MT and undergo a tug-of-war. Once resolved, the directional outcomes at 2D MT intersections have a straight to turn ratio of 1.1; whereas at 3D MT intersections, liposomes more frequently go straight (straight to turn ratio of 1.8), highlighting that spatial relationships at intersections bias directional outcomes. Using 3D super-resolution microscopy (STORM), we define the gap between intersecting MTs and the liposome azimuthal approach angle heading into the intersection. We develop an *in silico* model in which kinesin-1 motors diffuse on the liposome surface, simultaneously engage the intersecting MTs, generate forces and detach from MTs governed by the motors’ mechanochemical cycle, and undergo a tug-of-war with the winning team determining the directional outcome in 3D. The model predicts that 1-3 motors typically engage the MT, consistent with optical trapping measurements. Modeled liposomes also predominantly go straight through 3D intersections over a range of intersection gaps and liposome approach angles, even when obstructed by the crossing MT. Our observations and modeling offer mechanistic insights into how cells might tune the MT cytoskeleton, cargo, and motors to modulate cargo transport.

**Significance Statement:** Kinesin-1 molecular motors transport vesicles containing essential cellular resources along the dense 3D microtubule (MT) cytoskeleton, with dysfunctions linked to neurodegenerative diseases such as Alzheimer’s. Despite its importance, the mechanism by which kinesin-1s maneuver intracellular cargoes through MT-MT intersections towards their destination remains unclear. Therefore, we developed a 3D *in vitro* model transport system, which challenges kinesin-1 motor teams to maneuver vesicle-like liposomes through MT-MT intersections. Surprisingly, liposomes are biased to pass straight through 3D MT intersections rather than turn, even when the MT intersection presents as a physical barrier. A mechanistic model informs this observation, suggesting that spatial relationships between the cargo and MT intersection influence how molecular motors maneuver intracellular cargoes towards their destination to satisfy cellular demands.

## Introduction

Long-range, secretory vesicle transport is powered by teams of kinesin molecular motors along the cell’s microtubule (MT) cytoskeleton (1–4), a dense, 3-dimensional (3D) network with numerous MT-MT intersections (5–7). These MT intersections present physical obstacles through which kinesin motors must navigate their cargo. Therefore, defining the biophysical mechanisms that govern the directional outcome at a MT intersection is essential to understanding kinesin-based intracellular cargo transport and its regulation in general.

Studying cargo transport in cells has offered insights into the physical challenges kinesins encounter at 3D MT intersections. For example, melanophore cargoes bend the MTs that form the intersections (8), while lysosomes pause when the gap between intersecting MTs is smaller than the lysosome itself (i.e., <100 nm) (9). Additionally, individual cargoes appear to spin at MT intersections (10), while cargoes also undergo azimuthal motion around the MT on which they are moving as a means of maneuvering past an intersection (11). Collectively, these data suggest that MT intersections obstruct cargo transport before the cargo goes straight or turns at the intersection. To better define such complex cargo behavior, kinesin-based cargo transport has been reconstituted *in vitro* as a means of controlling the complexities inherent to cellular systems. Specifically, when kinesin ensembles were coupled to stiff bead cargoes and challenged with 2-dimensional (2D) MT intersections on a glass surface, cargoes paused at intersections, suggesting a tug-of-war between opposing groups of motors interacting with the intersecting MTs, prior to switching tracks or passing through (8, 12, 13). In contrast, directional outcomes for beads transported by kinesin-1 motors through 3D MT intersections exhibited a geometry-dependent bias towards passing through or switching to the intersecting MT, highlighting the importance of 3D spatial relations between the cargo and intersecting MTs (14, 15). Common to these studies is that kinesin motors were fixed to and immobile on the cargo surface as compared to *in vivo* vesicular cargoes, which have fluid-like membranes that allow motors to diffuse on the cargo surface. Therefore, investigators developed approaches to attach motors to lipid-bound vesicles (16, 17). Experimental (16, 18, 19) and theoretical work (20, 21) suggest that motors freely diffusing on the cargo surface can better navigate their cargoes past obstacles (19) and bind more efficiently to MTs, thus promoting longer run lengths compared to immobile motors fixed to the cargo surface (20).

Here, we build complexity *in vitro* by challenging fluid-like liposome cargoes (16, 22, 23), decorated with a controlled number (≤20) of bound kinesin-1 motors, to navigate either 2D or 3D MT intersections (Fig. 1, Movies S1-S3). Liposomes often pause at 2D and 3D MT intersections before continuing straight or turning with equal probability at a 2D MT intersection. Intriguingly, at a 3D MT intersection, liposomes predominantly go straight. To understand these directional outcomes, we developed a mechanistic model of kinesin-cargo transport both prior to and during an intersection encounter. The model predicts that directional outcomes in a 3D intersection occur following a tug-of-war between individual motor teams on the liposome surface where typically 1-3 motors per team engage each of the intersecting MTs. The bias to go straight through a 3D MT intersection highlights that spatial relationships between cargo and the intersecting MTs influence the cargo’s directional outcome and suggests that cells could modulate these spatial relationships to guide cargo towards its specific subcellular destination.

**Figure 1.**
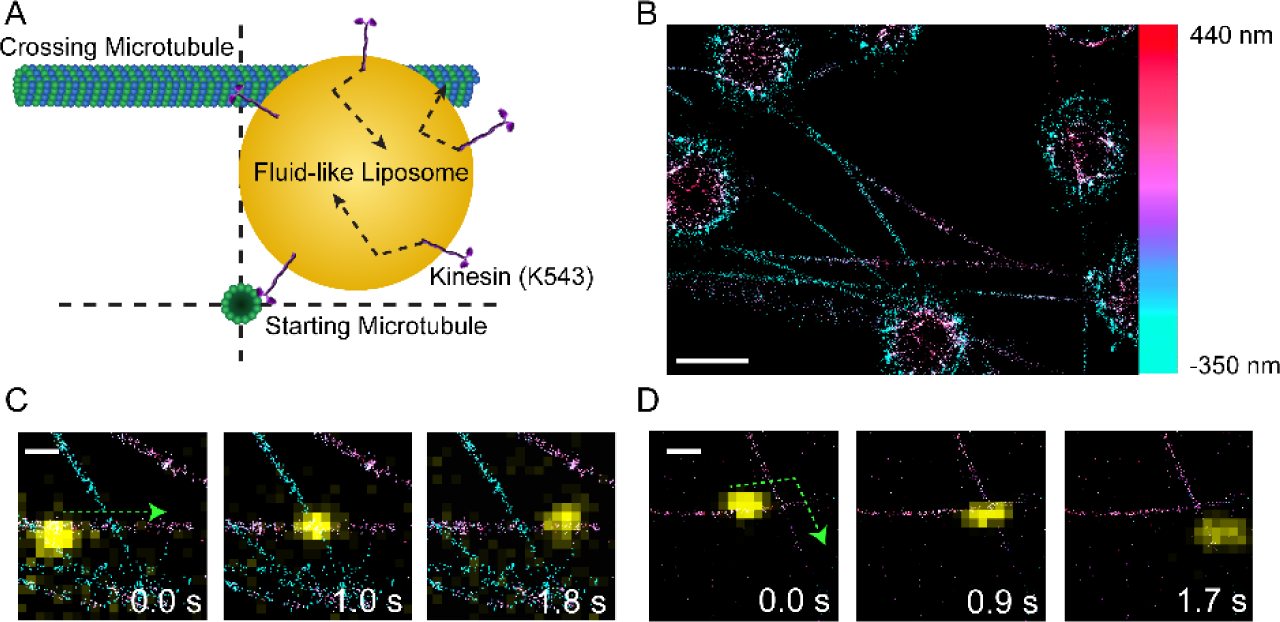
*3D* MT Intersection Model System. (A) Illustration of a liposome cargo (yellow sphere) transported by multiple kinesin-1 motors (purple) approaching an intersection between two MTs (green). Dashed lines indicate the plane which contains the MT intersection. (B) 3D STORM image of suspended MT intersections. Color bar represents Z-axis scale. Scale bar = 3 μm. (C) Movie frames of a liposome (yellow) proceeding straight through an MT intersection. Outcome shown by dashed green arrow. MT gap = 250 nm. Scale bar = 500 nm. (D) Movie frames of a liposome (yellow) turning in a MT intersection. Outcome shown by dashed green arrow. MT gap = 130 nm. Scale bar = 500 nm.

## Results

### Long distance liposome transport by kinesin-1 motor ensembles: evidence for more than one engaged motor

Fluid-like, 350 nm diameter liposomes, an *in vitro* model of membrane-bound vesicles, were transported by ensembles of ∼5-20 constitutively active, truncated kinesin-1 motors along single MTs attached to a glass coverslip (see Methods, Fig. S1). Liposome run length and velocity were measured using kymography and compared to Qdot-labeled single kinesin-1 molecules in saturating MgATP at 22° C (Fig. 2). Single kinesin-1 molecules step processively with a median run length of 1.3 μm (95% CI 1.2 – 1.6 μm) at a velocity of 890 ± 20 nm/s (mean ± SEM, Figs. 2B, S2), comparable to that reported previously (24). In contrast, liposomes transported by ensembles of ∼5 or more kinesin-1 motors (see Methods and Fig. S1 for motor counting), had median run lengths up to 12-fold longer than a single motor (Figs. 2B, S3). In fact, ensembles of ∼10 and ∼20 kinesin-1s per liposome had undefined upper 95% CIs (Figs. 2B, S3B), suggesting that run length was track-length-limited. Liposome velocity was similar across the ∼5, ∼10, and ∼20 kinesin-1 ensemble sizes (670 – 800 nm/s, Fig. S3A). However, these velocities were all slower than the single Qdot-labeled motor velocity (Fig. S2A). These data suggest that during ensemble transport more than one kinesin-1 motor may engage the microtubule simultaneously, which we confirmed by optical trapping measurements.

**Figure 2.**
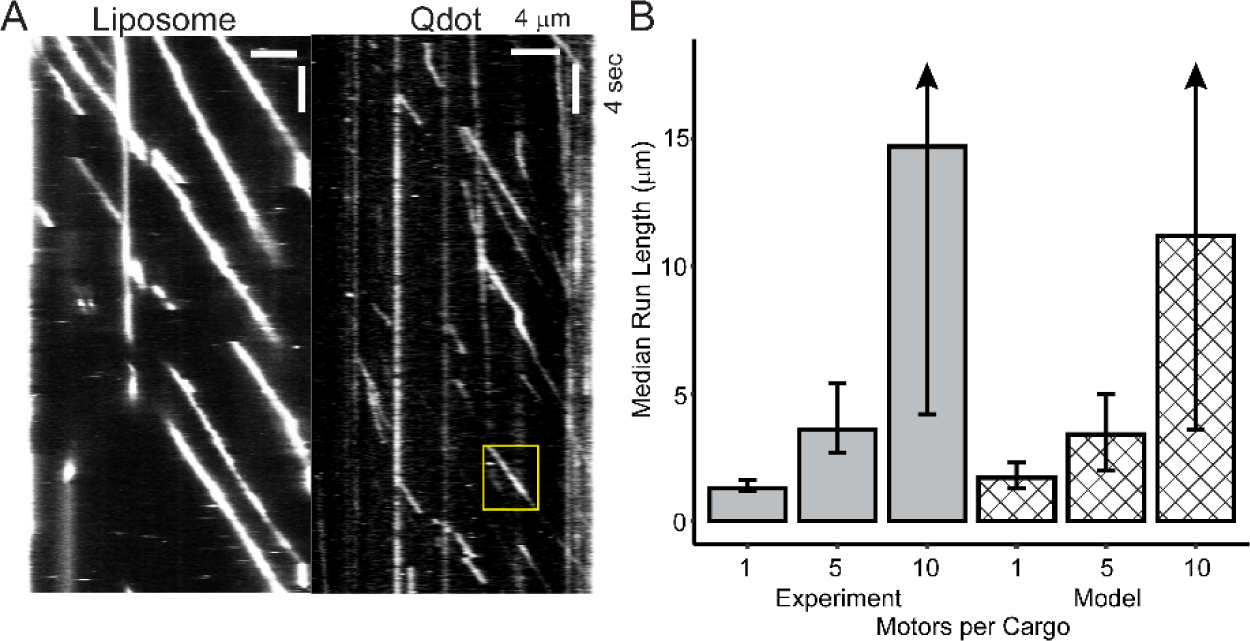
Multiple Kinesins Can Simultaneously Engage the MT. (A) Kymographs of cargo motility along a single MT in 2D. Left kymograph is liposome motility with 10-kinesins, right kymograph is single-kinesin Qdot motility. Vertical axis represents time and horizontal axis represents distance. Bounding box (yellow) indicates a single run by a kinesin-transported Qdot where the horizontal axis gives the run length, the vertical axis gives the event lifetime, and the slope of the streak gives the velocity. (B) Bar graph of median run length from experimental (grey) and modeled (crosshatch) kinesin-1 motility versus number of motors per cargo. Experimental N = 297, 161, and 137 events for 1, 5, and 10 motors per cargo, respectively. Error bars indicate upper and lower 95% CI. Arrow indicates that the bound could not be defined. Modeled trajectories are cut off after 15s, which gives a maximum run length of ∼11 μm.

For optical trapping purposes, 500 nm silica beads were coated with the same lipid mixture used to create the liposome cargoes (see Methods). Either limiting or a 20-fold molar excess of kinesin-1 motors were conjugated to the lipid-coated beads. At limiting kinesin-1 incubation, once a bead was captured in the optical trap and brought into contact with a MT (Fig. 3A), only ∼1 in 10 beads bound to the MT and produced force, suggesting that these MT-binding events were from beads conjugated to a single kinesin-1 motor. For these events, the kinesin-1 motor pulled the bead against the resistive trap force, which increased until the kinesin-1 underwent an abrupt force-dependent detachment from the MT, returning the bead to the trap center and thus baseline force (Fig. 3B inset). The force on the bead just prior to detachment was well-described by a Gaussian distribution centered at 5.7 ± 1.1 pN (Fig. 3C), similar to that reported for single kinesin-1 motors (25–27). For multi-motor, lipid-coated beads, a 20 kinesin-1 motors per bead incubation ratio was chosen so that the surface density of kinesin-1 motors was equivalent to that of a 350 nm liposome with ∼10 bound kinesin-1. Force ramps for these multi-motor beads (Fig. 3B) could reach up to 3-fold higher forces, confirming the presence of multiple engaged motors. The force prior to detachment showed a broader distribution (Fig. 3C), which was best described by the sum of three Gaussians centered at 5.8 (Amplitude = 0.7), 11.1 (Amplitude = 0.17), and 15.4 pN (Amplitude = 0.13). Log-likelihood ratio testing showed that inclusion of the third Gaussian is statistically justifiable (P = 5 x 10^-6^). Interestingly, 70% of the events were in the peak centered at 5.8 pN, suggesting that most force ramp events for a motor ensemble have only a single engaged kinesin-1 motor. The remaining 30% of events are distributed in the higher force peaks, suggesting less frequent engagement of 2 or 3 motors from the ensemble. The median event lifetime for single motor force ramps was 1.6 s while multi-motor force ramps had a median event lifetime of 1.9 s (Fig. S4), only a modest increase over the single motor condition. While these similar lifetime distributions are further evidence that multi-motor liposomes predominantly have a single kinesin-1 motor engaged with the MT, the higher detachment forces confirm that multiple motors can simultaneously engage the MT.

**Figure 3.**
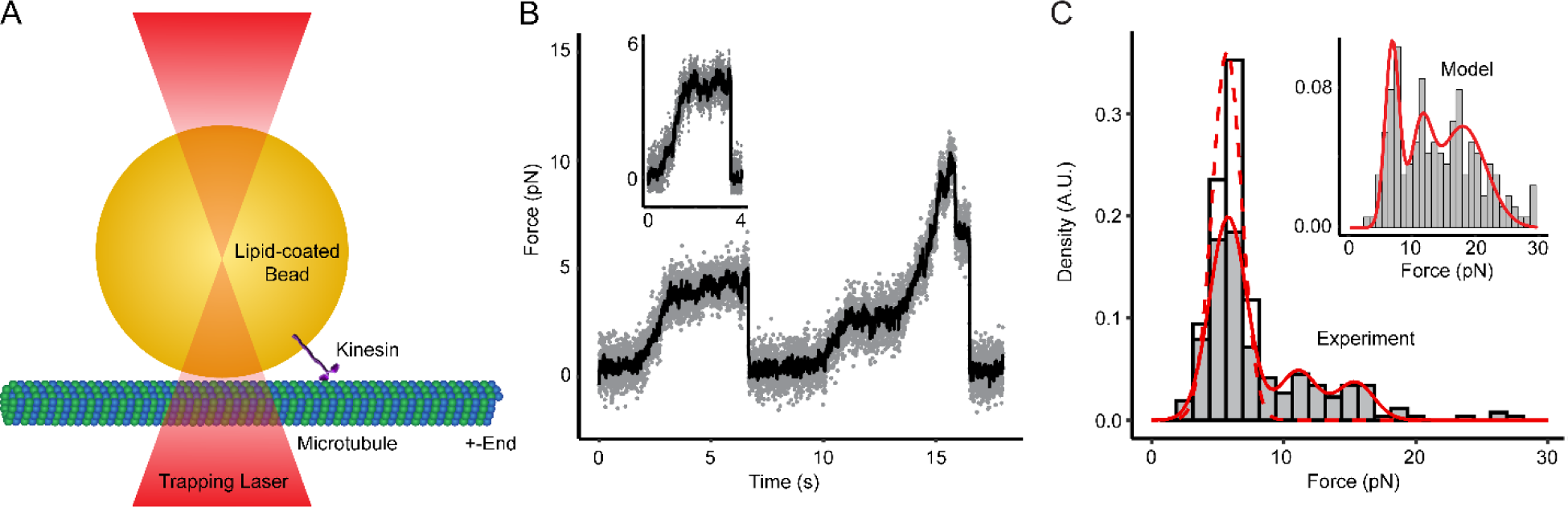
Multiple Kinesins Can Generate High Forces in an Optical Trap. (A) Schematic of optical trap ramp force assay. (B) Example optical trap force trace for a lipid-coated bead transported by multiple kinesin-1 molecules. Inset, example force trace for a lipid-coated bead transported by a single kinesin-1 molecule. (C) Histogram of maximum force prior to detachment for beads incubated with 20-fold molar excess of kinesin-1 (grey bins) and single kinesin-1 transported beads (white bins). Red solid curve shows triple Gaussian fit to the multi-motor data (A_1_ = 0.70, A_2_ = 0.17, μ_1_ = 5.8 pN, μ_2_ = 11.1 pN, μ_3_ = 15.4 pN). N = 213 events from 5 independent bead preparations. Dashed red curve shows Gaussian fit of maximum force prior to detachment for single kinesin-1 transported bead (μ = 5.7 pN, σ = 1.1 pN). N = 34 events from 3 independent bead preparations. Inset, histogram of modeled maximum force prior to detachment for kinesin-1 ensembles. Red curve shows triple Gaussian fit to the data (A_1_ = 0.33, A_2_ = 0.18, μ_1_ = 6.8 pN, μ_2_ = 11.7 pN, μ_3_ = 17.8 pN). N = 202 simulated trajectories.

### Directional outcomes of liposome-kinesin-1 complexes at 2D MT intersections depend on ensemble size

To create simple 2D MT intersections *in vitro*, we adhered MTs to a coverslip surface. By differential MT fluorescence intensity labeling, we could identify which MT was on top versus the bottom of the intersection (see Methods, Fig. 4A, Movie S3). Knowing this, we determined whether liposomes approached an intersection from either the bottom or top MT. For bottom MT approaches, the crossing MT presents as an obstacle the liposome must overcome, while a liposome approaching on the top MT may avoid the bottom crossing MT. For either case, we recorded the directional outcomes at the intersection (i.e., straight, turn, or terminate) for liposomes transported by ∼5, 10, or 20 kinesin-1 motors. For bottom MT approaches (Fig. 4B), liposomes with ∼5 kinesin-1s were more likely to go straight (56%) than to turn (33%) with infrequent terminations (11%). In contrast, liposomes with ∼10 or ∼20 kinesin-1s went straight and turned with nearly 50% probabilities, terminating infrequently (Fig. 4B). For top MT approaches (Fig. 4C), liposomes with approximately 5, 10, or 20 kinesin-1s rarely terminated and went straight in ≥75% of events with the remainder of events turning at the intersection. The observed high likelihood of going straight when traveling on the top MT suggests that the bottom MT does not present as a physical barrier. However, the fact that liposomes can turn onto the bottom MT suggests that in some cases the bottom MT is within reach of motors on the liposome surface.

**Figure 4.**
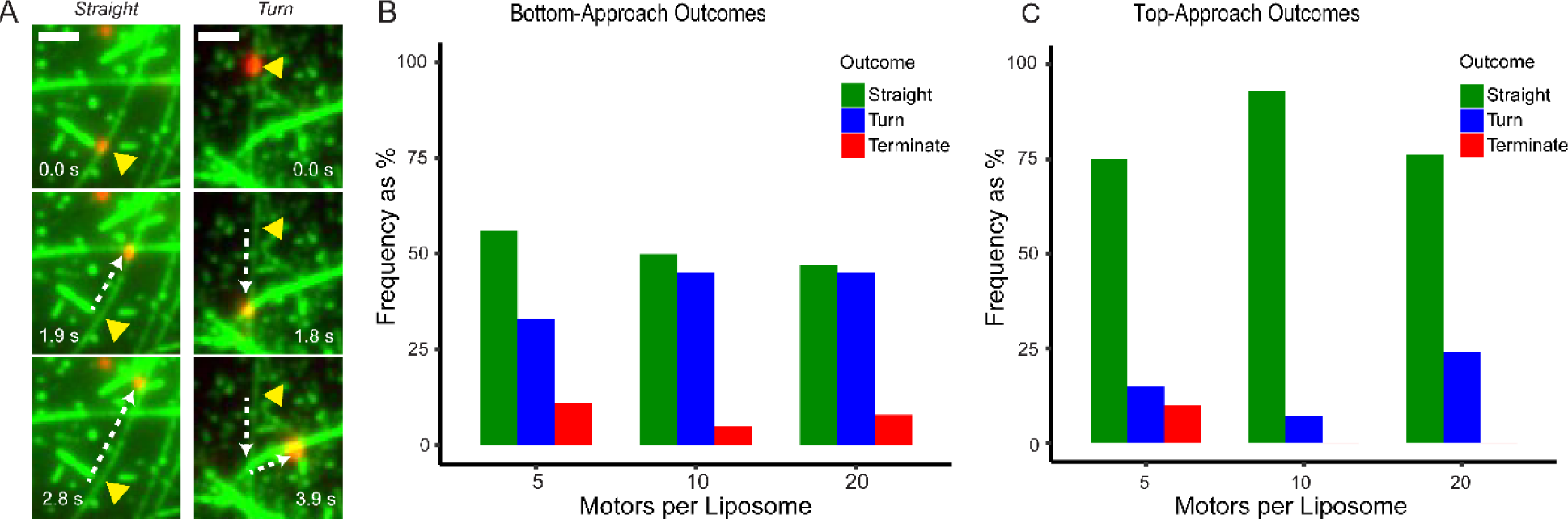
Directional outcomes for kinesin-liposome complexes in 2D MT intersections depend on the approach MT and motor number. (A) Timestamp images of a 10-kinesin-liposome complex passing straight (left) or turning (right) in a 2D MT intersection. In both cases, the liposome approaches the MT intersection on the dimly-fluorescent, bottom MT (see Methods). Yellow arrowhead indicates starting liposome position. (B) Bar graph of directional outcomes for liposomes approaching the intersection on the bottom MT. N = 50, 38, and 60 events for 5, 10, and 20 motors per liposome, respectively. Data from at least 3 separate liposome preparations. (C) Bar graph of directional outcomes for liposomes approaching the intersection on the top MT. N = 44, 42, and 41 events for 5, 10, and 20 motors per liposome, respectively. Data from at least 3 separate liposome preparations.

Kymographic analysis revealed that liposomes transported by approximately 5, 10, or 20 motors when approaching the intersection on the bottom MT pause at intersections with increasing frequency (26%, 42% and 67%) and median lifetimes (1.5s, 2.7s, and 3.7s), respectively (Fig. S5A). For top MT approaches (Fig. S5B), liposomes paused less frequently on average (18%, 17%, and 44%) but with similar median lifetimes (1.2s, 1.1s, and 3.3s) for all motor numbers. Pausing suggests that kinesin-1 motor teams simultaneously engage both the top and bottom MT of an intersection and undergo a tug-of-war, which must be resolved before a directional outcome is determined.

### Liposome transport on MT tightropes

Although directional outcomes through 2D MT intersections offer insights into motor ensemble navigation at an intersection, intracellular vesicle transport occurs within the 3D cellular environment. Thus, 3D MT intersections are a more physiologically relevant challenge. However, we first characterized liposome transport along a suspended MT where the motor ensemble has 360° access to the MT surface. Prior work has shown that single kinesin-1 molecules predominantly follow a MT protofilament (28–30), leading to a mix of straight and helical trajectories because *in vitro* MTs have variable protofilament numbers and consequently super-twisted protofilaments (30). In fact, liposomes transported by ∼10 kinesin-1 motors show a similar mixed trajectory pattern along single suspended MTs (see Methods for analysis). Specifically, 35% of trajectories were best fit by a straight trajectory, while 65% were well-fit by a 3D sinusoid (Figs. 5A, S6), indicative of a helical trajectory. The absolute magnitude of the helical pitch ranged from 3 to 10 μm per turn, with roughly equal fractions of helical trajectories being left-versus right-handed (Fig. 5B). This trajectory distribution is similar to that reported previously for a single kinesin-1 motor (30), suggesting that liposomes transported by kinesin-1 ensembles follow MT protofilaments.

**Figure 5.**
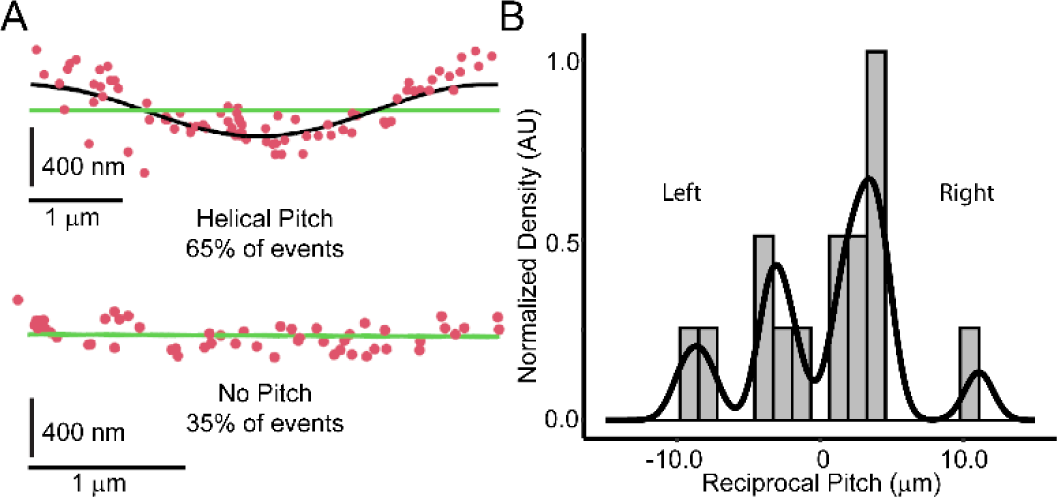
Liposomes can take helical paths on suspended MTs. (A) Top, example of helical liposome trajectory. Green line represents MT, red dots represent liposome position along the MT, and black curve represents 3D sinusoidal fit (see Methods, Fig. S5). Bottom, example of straight liposome trajectory. Horizontal scale bar refers to position along the MT, vertical scale bar refers to liposome center vertical position relative to MT center. (B) Histogram of reciprocal pitches measured in helical liposome trajectories. Black curve is the kernel density function and is for visualization purposes only. N = 23 total events, 15 helical events.

### Directional outcomes of liposome-kinesin-1 complexes at 3D MT intersections depend on MT-MT gap and liposome approach angle

To create 3D MT intersections, MTs were flowed through perpendicular channels of a flowcell having poly-L-lysine-coated silica beads of mixed diameter (range of 1.0 – 3.0 μm) on the flowcell surface as pedestals from which MTs were suspended. This procedure resulted in nearly perpendicular 3D MT intersections with a variable gap between MTs at the point of intersection (Fig. 1B-D). For these experiments, we limited our study to 350 nm liposomes transported by ∼10 kinesin-1 motors.

Since liposomes can travel in a helical trajectory while approaching the intersection (see above), a liposome may or may not interact with the intersecting MT depending on the liposome approach angle, α. As a point of reference, an α of 0° describes a liposome with its center vertically above the MT it is being transported on, which is the same side the intersecting MT is on (Fig. 6A). Based on the intersection gap (*d*) and the liposome approach angle (α) (Fig. 6A), liposome-intersection encounters were split into two classes; those where kinesin-1s on the liposomes could reach the crossing MT, and those where the kinesin-1s could not (Fig. 6B). The directional outcomes, for *d* and α pairs that permitted interactions between kinesin-1s on the liposome and the crossing MT, had a higher probability of going straight through an intersection (57%) compared to turning (31%) or terminating (12%) (Fig. 6C). As in 2D intersections, liposomes often paused in the intersections (70% of events) with a median lifetime of 0.75s (Fig. S7A and S7B). Pausing was less frequent in events that went straight (59%) than in events that turned (93%). Pausing at 3D intersections suggests that motor teams on the starting and crossing MT engage in a tug-of-war, which must resolve prior to the liposome either going straight or turning. However, the ∼2-fold propensity to go straight in a 3D intersection (i.e., a straight to turn ratio of 1.8) compared to a 2D intersection (ratio of 1.1) for the same ∼1:10 liposome-kinesin-1 complex is unique to the 3D MT intersections.

**Figure 6.**
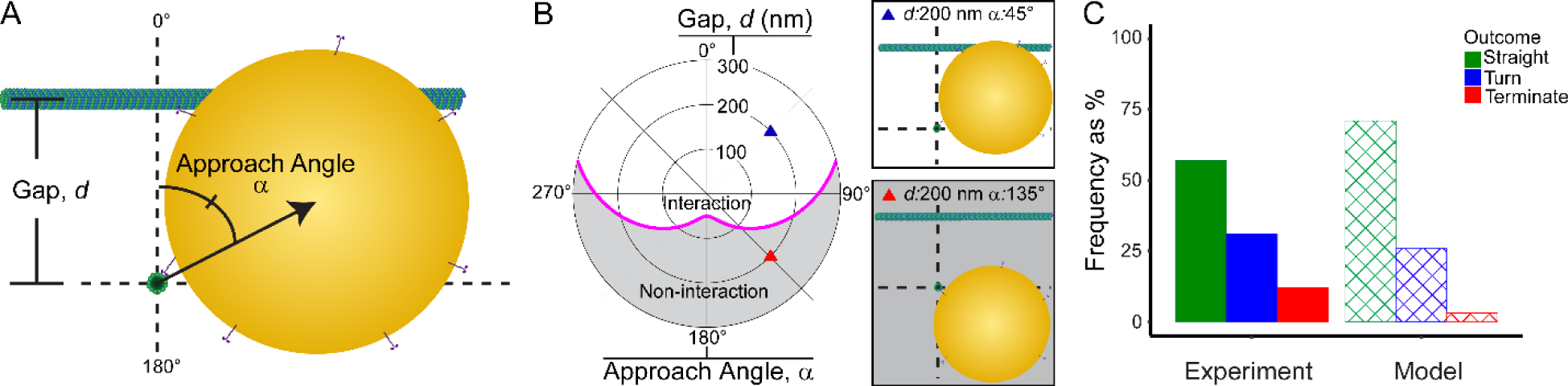
Kinesin-liposome complexes preferentially go straight in 3D MT intersections. (A) To-scale schematic of a kinesin-liposome complex approaching (into the image plane) a 3D MT intersection. Two geometric parameters describe the spatial relations between the liposome and the intersection, the Gap (*d*) between the MTs along the Z-axis and the Approach Angle (α), which describes the position of the liposome relative the crossing MT where 0° is up towards and 180° is down away from the crossing MT. (B) Left: Polar plot of (*d*, α). Magenta line divides interaction and non-interaction geometries for liposome with intersecting MT. Right Top: To-scale schematic of an interaction geometry. Right Bottom: To-scale schematic of a non-interaction geometry. (C) Bar chart of directional outcomes in 3D MT intersections for liposome-MT interaction events only, experimental (solid, N = 100 events) and modeled (crosshatch, N = 178 simulations).

Knowing the precise spatial relationship between the liposome and the MTs within the intersection, we determined if these spatial relationships biased directional outcomes. Therefore, we plotted directional outcomes as a function of intersection gap (*d*) and liposome approach angle (α) on a polar scatter plot, color coded by outcome (Fig. S8). As expected, 100% of liposome-kinesin-1 complexes that could not physically interact with the crossing MT (Fig. S8, below magenta curve) passed straight through the intersection. In contrast, all possible directional outcomes were observed for liposome-kinesin-1 complexes which could interact with the crossing MT (Fig. S8, above magenta curve). To identify spatial relationships where certain directional outcomes may be enriched, we transformed the data into a heatmap with 4 spatial quadrants color coded by a straight to turn ratio (Fig. 7, Experiment). Based on this ratio, going straight through an intersection was more common than turning (i.e. straight to turn ratio >1) for all quadrants, consistent with the overall directional outcomes data (Fig. 6C). However, subtle differences in the straight to turn ratio do exist with the highest probability of going straight when α > 75° and *d* > 175 nm, whereas turning occurred with the highest relative probability when α > 75° and *d* <175 nm (Fig. 7, Experiment). Thus, 3D spatial relations between the liposome and the intersecting MTs do influence the directional outcomes.

**Figure 7.**
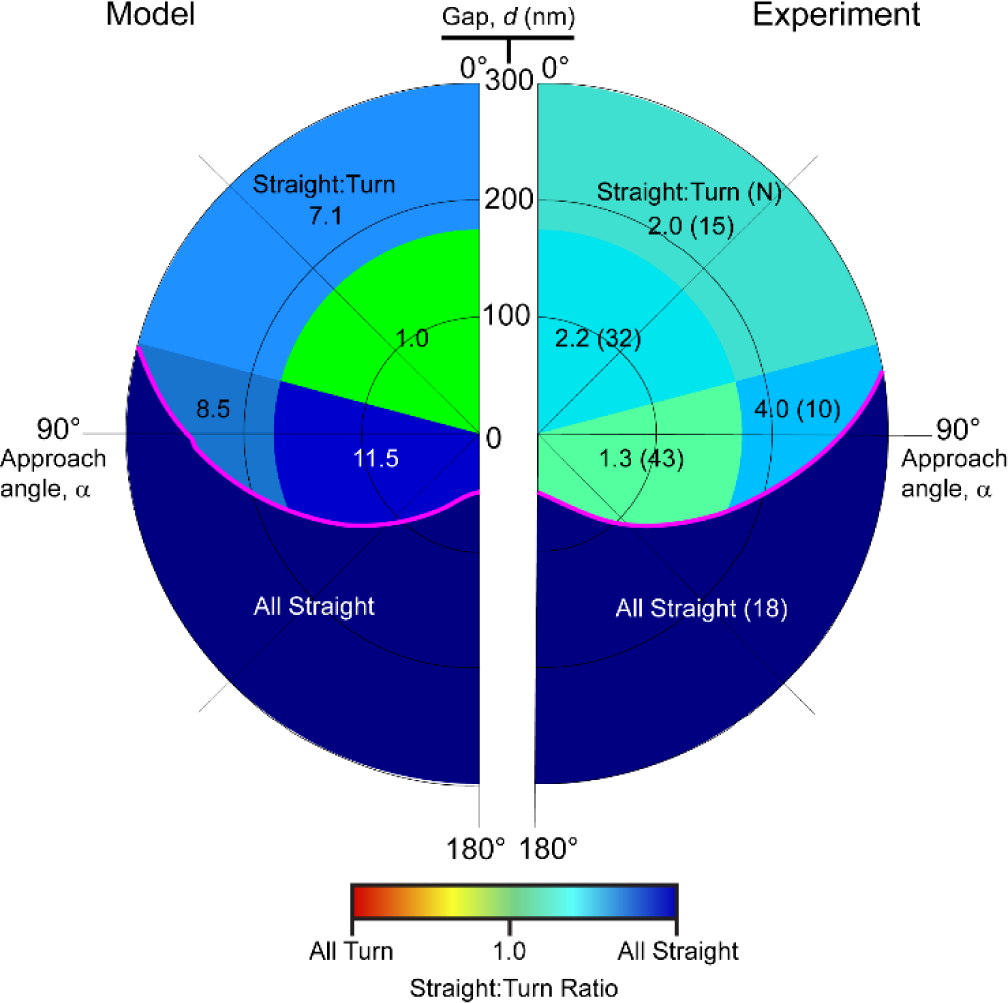
3D directional outcomes depend on spatial relations between the liposome and the intersecting MTs. Heatmap of modeled (left) and experimental (right) straight to turn ratios as a function of approach angle (α) and gap (*d*). Events are split into 4 zones based on MT gap (>175 nm or <175 nm, i.e., the liposome radius) and approach angle (>75° or <75°), with the fifth zone being non-interaction events. Because termination is rare, only straight to turn ratio is shown. Straight to turn ratio is color-coded (see color bar). Total N = 118 events for experimental data set, 100 interactions and 18 non-interactions. For modeled events, N = 210 events total, 178 interactions and 32 non-interactions.

### Mechanistic model of liposome transport by ensembles of kinesin-1 motors

To mechanistically understand the behavior of a liposome-kinesin-1 complex in a 3D MT intersection, we developed an *in silico* model of liposome-kinesin-1 complex transport (see Supplementary Information for model details). The model: 1) incorporates the mechanochemical cycle of kinesin-1 as defined previously (31) with some modifications (see Supplementary Information); 2) explicitly simulates diffusion of kinesin-1s on the liposome surface; 3) simulates the translational and rotational diffusion of the liposome, and; 4) calculates the forces generated by and applied to each individual kinesin-1 motor. Knowing the forces on each attached kinesin-1 motor, the model predicts the motor’s stepping activity and detachment probability and, with these, the 3D trajectory of the liposome.

Using this model, we simulated liposome motility along single microtubules and measured the run lengths for liposomes with 5 and 10 kinesin-1s, which exceeded the simulated single molecule run lengths, as experimentally observed (Fig. 2B). Additionally, regardless of the kinesin-1 motor number, liposome velocity was similar at ∼800 nm/s as observed experimentally (see Supplementary Information, Fig. S3). Thus, the model recapitulates multi-motor liposome transport in the absence of external forces.

To determine whether the model successfully predicts the number of engaged motors under external load, we simulated liposome motility against an external load from an optical trap and measured the detachment forces and event lifetimes for liposomes with 1 or 20 bound kinesin-1 motors (Movie S4 and S5). For a liposome with a single kinesin-1, the model predicts a mean detachment force of 7.3 ± 2.4 pN (mean ± SD, N = 100 simulations). For the 20 kinesin-1 case, the model predicts detachment forces distributed as the sum of three Gaussians with peaks centered at 7.7, 14.2, and 20.7 pN (Fig. 3C, inset) with a median event lifetime of 2.9 s (Fig. S4, dotted lines). This result is consistent with our experimental observation of 3 peaks of detachment forces which were roughly multiples of the lowest force peak, and the lowest force peak being similar to the average force of a single motor. The amplitude of the lowest force peak in both the experiment (A = 0.70) and the model (A = 0.42) was the greatest, suggesting qualitative agreement between the model and experiment that most often only a single transporting motor is engaged and >3 engaged motors was extremely rare.

Based on the agreement between the modeled and experimental results above, we then simulated liposome transport by 10 kinesin-1 motors through 3D MT intersections (see Supplementary Information for modeling details). The simulated overall directional outcomes are in good agreement with the experimental results (Fig. 6C). Specifically, liposomes are predicted to pass straight through an intersection (71%) more often than turn (26%), with termination being rare (3%) (Fig. 6C). Furthermore, the model predicts pausing behavior in intersections, with 51% of events having a measurable pause with a median lifetime of 1.0s (Fig. S7). Pausing was more common in events that turned (100%) than in events that went straight (31%). These predicted directional outcomes and pausing statistics agree qualitatively with our experimental observations.

As in the experiment, the precise geometry of the liposome relative to the intersecting MTs is known within the model. Therefore, we created a polar heatmap of simulated 3D intersection directional outcomes, described as straight to turn ratios distributed into quadrants (Fig. 7, Model). Much like the experimental results, all 4 quadrants had a straight to turn ratio ≥1, predicting that most spatial relationships bias liposomes toward going straight through the MT intersection (Fig. 6C), even though apparent quantitative differences between the model and experiments do exist.

## Discussion

Long-range kinesin-1-based vesicular transport along the cell’s complex, 3D microtubule network requires motors to maneuver their cargo through numerous MT intersections (5). To define the underlying mechanism by which a specific directional outcome occurs at a MT intersection, we created an *in vitro* model system in which fluid-like liposome cargoes decorated with kinesin-1 motors encountered MT intersections. Interestingly, even though motors on the liposome surface interact with the intersecting MT, liposomes predominantly go straight through 3D MT intersections. Our development of an *in silico* mechanistic model provides insight into how this is possible when the intersecting MT creates a physical barrier.

### Spatial relationships matter for directional outcomes at MT intersections *in vitro*

The most striking example of how liposome directional outcomes are biased by the spatial relationship between the liposome and the MT intersection is evident when comparing outcomes for 2D versus 3D MT intersections. Specifically, for 2D intersections, liposomes with ≥10 kinesin-1 motors approaching the intersection on the bottom MT show no directional bias, going straight and turning equally (Fig. 4B). Whereas in 3D MT intersections, liposomes transported by ∼10 motors show a clear preference for going straight, as successfully modeled *in silico* (Fig. 6C). The most obvious spatial difference between 2D and 3D MT intersections that may contribute to this difference is MT access. In 3D, liposomes have 360° access to the MT surface that they’re traveling on when approaching the intersection (Figs. 5, 6A, 6B), whereas in 2D, <180° of the MT surface is accessible due to the slide surface. Thus, liposomes can simply avoid the crossing MT in 3D. However, even if the liposome can’t avoid the intersecting MT, liposomes predominantly go straight through 3D MT intersections rather than turning or terminating (Figs. 6C, 7). This directional outcome bias is in contrast to a study where 1 μm silica bead cargoes, transported by ∼30 kinesin-1 motors fixed to the bead surface, preferred to turn when the MT gap for a 3D MT intersection was ≤0.5 μm, only preferring to go straight when the MT gap is greater than or equal to the cargo diameter (15). This disparity suggests that the liposome’s fluid-like phospholipid membrane, which allows motors to diffuse on the liposome surface, contributes to biasing liposomes to go straight at a 3D MT intersection. In addition, the predicted ability of the fluid-like liposome to diffusively spin on its axis while being transported or at a 3D MT intersection (see Movies S6-S10), as reported in cells (10), may also contribute to the directional bias at 3D MT intersections.

### MT intersection directional outcomes result from motor teams engaged in a tug-of-war

Liposomes frequently pause at both 2D and 3D MT intersections (Results; Figs. S5 and S7) which, in our model, results from small individual teams of kinesin-1s on the liposome surface being simultaneously engaged with both of the intersecting MTs and thus undergoing a tug-of-war, as suggested previously (12, 13, 15, 22). Since the number of engaged motors in either team is stochastic and generally varies between 1-3 motors, it’s this dynamic motor binding and unbinding within a team (32) (Figs. 3B and 3C) that contributes to the eventual directional outcome, which is simply dictated by the motor team that wins the tug-of-war (Fig. 8, Movies S6 and S7). Similarly, in cells, cargoes have been reported to pause at MT intersections before turning or proceeding straight, suggesting that a tug-of-war is at play (8–10).

**Figure 8.**
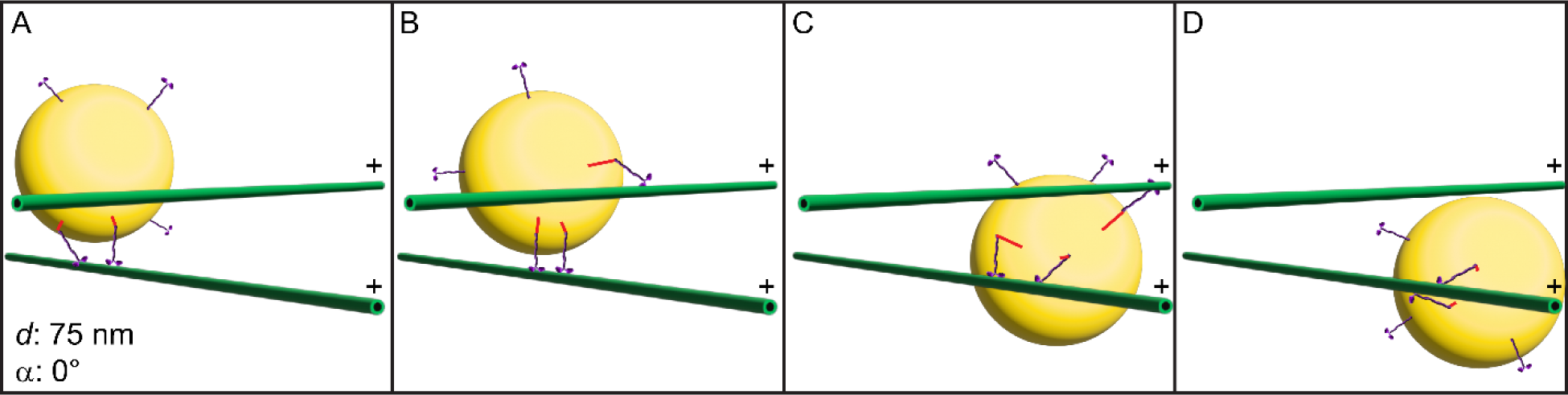
Model-simulated liposome proceeds straight through a MT intersection following a tug of war between motors engaged with each of the intersecting MTs. In all frames: liposome (yellow sphere), kinesin (two heads with purple tail connected to liposome with red line representing the magnitude and direction of force on the corresponding kinesin), MTs (green rods with +-end indicated). (A) Liposome approaches (α = 0°) an intersection (*d* = 75 nm). (B) A new kinesin engages the crossing MT, entering into a tug of war with the kinesins engaged with the starting MT. (C) The force of the kinesin attached to the crossing MT rotates the liposome around the starting MT and thus underneath the crossing MT. (D) The kinesin on the crossing MT detaches, allowing the liposome to proceed straight through the intersection.

Because straight outcomes are preferred in 3D MT intersections, there must be other mechanism(s), in addition to the tug-of-war, that bias the directional outcome. For example, model simulations suggest that a liposome undergoes significant angular wobble (i.e., Brownian motion), which allows a new motor to attach to a different protofilament on the original MT. Once this motor binds, the liposome’s center of mass reorients above the new motor team, favorably positioning the liposome to pass beneath the crossing MT (Movie S8 and S9). Another example of liposome reorientation contributing to a straight outcome occurs with small intersection gaps when a motor engages the crossing MT, effectively rotating the liposome so that when the motor on the crossing MT detaches, the liposome continues straight (Fig. 8, Movie S10). Angular cargo displacements at MT intersections in cells have been reported (11), emphasizing that such displacements, whether diffusional or motor-generated, may be a key mechanism for maneuvering cargoes through dense MT networks. Subtle differences between the modeled and experimental outcomes as shown on the polar heatmap (Fig. 7) suggest additional mechanisms not captured by the model may bias outcomes. For instance, forces applied to the kinesin motor that are normal to the MT may also promote detachment (33) and could explain spatial arrangements where more turning is observed in the experiment than in the model (Fig. 7; see Supplementary Information for more detail). Furthermore, kinesin-induced changes to the MT lattice spacing, which promote kinesin binding (34–38), may also play an important role.

### Kinesin ensembles are anti-cooperative on a fluid-like liposome cargo

Understanding the nature of motor-motor interactions on a physiologically relevant, lipid-based cargo is key to understanding the dynamics of a tug-of-war in an intersection and how this might introduce bias in the directional outcome distribution. Prior studies with kinesin ensembles attached to rigid cargoes have reported anti-cooperativity, where binding of the first kinesin negatively influences binding of the next, perhaps due to mechanical interference between stiffly coupled motors (39–42). However, recent modeling efforts predict that kinesins bound to a lipid-based cargo, where motors can diffuse on the cargo surface, would behave cooperatively, i.e., MT engagement of the first kinesin promotes engagement of additional kinesins (20, 21). Based on our optical trapping experiments (Fig. 3B, C), rarely did more than 3 kinesins out of ∼20 on the fluid-like lipid-coated bead surface simultaneously engage the MT. In fact, most often only a single kinesin was engaged (Fig. 3C), as predicted by our model, suggesting anti-cooperative binding behavior. Why models of lipid-bound cargo transport predict different kinesin-MT binding behavior may lie in how the kinesin tail stiffness is represented.

Specifically, we model the kinesin tail as a Hookean spring with a set rest length that only allows the kinesin motor to bind the MT when the liposome is optimally positioned at a specific distance from the MT. This is in contrast to other models (20, 21, 32) which allow binding of the kinesin to the MT at any distance less than or equal to the length of the kinesin tail, i.e., a highly flexible or compressible tail. In our model, once the first motor binds, it restricts the spatial freedom of the liposome, making it more difficult for additional motors to bind the MT, providing a potential reason for anti-cooperative kinesin binding, similar to prior observations with myosin-Va (22, 23).

### Cargo sorting and polarized transport in cells through the MT cytoskeleton

Cells often utilize different motors to move different cargoes. Specifically, the kinesin-1, - 2, and -3 subfamilies are the primary drivers of MT plus-end-directed intracellular vesicular transport (2, 43, 44). Although motors from these families evolved to perform similar tasks and share a largely conserved motor domain, there are subfamily-specific differences in their mechanochemistry. Compared to kinesin-1 motors, kinesin-2s show more rapid force-dependent MT detachment (31, 45–47), while kinesin-3s have rapid kinetics of MT association (47–50). Does motor mechanochemistry lead to different directional outcomes in MT intersections? To address this question, we performed simulations varying only the rates of motor attachment to and detachment from the MT (see Supplementary Information). In fact, directional outcomes in a 3D MT intersection can be altered by varying these rates and no other parameters, suggesting that *in vivo* MT-based cargo transport and delivery may be matched to the cell’s physiological demands by fine-tuning the complement of kinesin family motors on the cargo.

In cells, cargoes are often sorted and moved in a polarized manner to specific subcellular compartments or cell surface locations (51–53). Given the density of the intracellular MT network, polarized cargo transport requires cargoes to be steered towards or away from specific subsets of MTs (51–53). Since our findings suggest that liposomes are biased to go straight through all 3D MT intersections, though the extent of this bias is tuned by spatial relationships at the intersection, there must be other intracellular regulatory processes to promote turning in 3D MT intersections, thus guaranteeing polarized cargo transport and sorting. For example, microtubule-associates proteins (MAPs) can alter the MT landscape in cells, such as MAP7 that promotes kinesin-1 binding to MTs (54–56), or Tau, which inhibits kinesin-1 binding (19, 41, 55, 57, 58). In addition, tubulin post-translational modifications such as acetylation and detyrosination, can alter kinesin-MT interactions directly or indirectly through changes in the MT lattice spacing (59, 60). Therefore, the cell possesses a significant and varied toolbox to potentially modulate directional outcomes at each MT intersection to help sort and polarize cargo transport in the cell’s complex 3D MT cytoskeleton.

## Methods

### Liposome-kinesin-1 complex preparation and characterization

Liposomes were prepared as described previously (16, 22, 23). In brief, a lipid mixture composed of (molar ratio) 84 parts DOPC (Avanti Polar Lipids), 5 parts PEG-ylated phospholipid (Avanti Polar Lipids), 5 parts cholesterol (Avanti Polar Lipids), 5 parts MBP:PE (Avanti Polar Lipids), and 1 part of the lipophilic dye, DiI (Thermo Fisher Scientific), was dried under nitrogen followed by 1 hour in vacuum (Rotovap;Eppendorf), then rehydrated overnight using PBS (pH = 7.4). Liposomes were extruded through a 1 μm pore diameter extruder (T&T Scientific) and incubated overnight with 1 μM thiolated neutravidin (SH-NaV). Excess SH-NaV was removed by pelleting the liposomes at 392,000 x g for 10 minutes and resuspending in PBS (pH = 7.4), repeated 3 times. Liposomes were then extruded to their final size of 350 nm diameter (22) through a 200 nm pore diameter extruder (T&T Scientific).

A truncated (aa 1-543), dimeric kinesin-1 (Mm kif5b) construct (K543) with a C-terminal FLAG tag and C-terminal biotin ligase recognition sequence was expressed using a baculovirus/Sf9 cell system (61). Functional kinesin dimers were purified via affinity chromatography to the C-terminal FLAG tag. An additional dimeric kinesin-1 for motor counting purposes on the liposome surface which had an N-terminal YFP, C-terminal FLAG tag and C-terminal biotin ligase recognition sequence was expressed and purified as above. Kinesin-1 motors were attached to liposome cargoes via the C-terminal biotin to the SH-NaV on the liposome surface. To create kinesin-liposome complexes of varying stoichiometries, kinesin-1 (4 μl of 250 nM to 1 μM dimer) was incubated with liposomes (20 μl of 10 nM) and 16 μl of Buffer 1 (25 mM Imidazole pH = 7.4, 25 mM KCl, 4 mM EGTA, 4 mM MgCl2) on ice for 1 hour.

To estimate the number of kinesin-1 dimers bound per liposome, we use a photobleaching approach described previously (22, 62). Liposomes were prepared without the lipophilic DiI fluorescent dye to minimize background fluorescence and incubated with varying concentrations of YFP-kinesin-1 to form kinesin-liposome complexes of varying stoichiometries. To promote landing of the liposomes on a microscope slide surface, slides were prepared with unlabeled microtubules attached to the slide surface via an anti-tubulin antibody (BioRad YL1/2). Kinesin-liposome complexes were added to the flow cell and imaged in TIRF. As the kinesin-liposome complexes landed on the slide surface, both the initial integrated YFP fluorescence intensity and the intensity decay over time were measured as the YFP molecules on the YFP-kinesins photobleached (Fig. S1A). The intensity per fluorophore was calculated as described in Nayak and Ruttenberg. Based on the initial intensity, the number of bound kinesins per liposome could then be estimated. Performing this analysis across a range of incubation stoichiometries yielded a standard curve (Fig. S1B) which was used to estimate the mean number of bound kinesins based on incubation conditions.

### Microtubule preparation

A mixture of unlabeled porcine brain tubulin (Cytoskeleton, Inc. Cat. No. T240) was copolymerized with either rhodamine-labelled tubulin (Cytoskeleton, Inc. Cat. No. TL590M) or Alexa-647-labelled tubulin (PurSolutions Cat. No. 064705). 40 μl of unlabeled tubulin at 5 mg/ml in BRB-80 buffer (80 mM PIPES pH = 6.9, 1 mM MgCl2, 1 mM EGTA) with 1 mM GTP was mixed with 10 μl of either rhodamine tubulin (2 mg/ml) or Alexa-647 tubulin (10 mg/ml) in BRB-80 buffer with 1 mM GTP. For differentially labelled bright and dim rhodamine MTs, dim MTs were made with 10 μl of 0.4 mg/ml rhodamine tubulin and bright MTs were made with 10 μl of 1.6 mg/ml rhodamine tubulin. The tubulin mixture was incubated on ice for 5 minutes, then centrifuged at 392,000 x g for 10 minutes before the supernatant was transferred to a fresh tube. The tubulin supernatant was incubated at 37° C for 20 minutes, prior to addition of paclitaxel to a final concentration of 20 μM prior to incubation at 37° C for an additional 20 minutes. The tube of assembled MTs was wrapped in foil to protect from photodamage and stored at room temperature.

### Qdot and liposome transport by kinesin-1 on single microtubules

For single motor motility assays, 2 μl of 1 μM Qdot 655 Streptavidin Conjugate (ThermoFisher) was incubated with 2 μl of 100 nM biotinylated kinesin-543 and 6 μl of Buffer 1 on ice for 1 hour prior to experiments. For ensemble motility assays, kinesin-liposome complexes were prepared as described above. Motility chambers were created by affixing a 22×22 mm no. 1 cover slip to a silanized 24×60 mm no. 1 cover glass with UV-curable adhesive and 125 μm Mylar shims. The motility surface was coated with 0.8% anti-tubulin antibody (BioRad YL1/2) in BRB-80 buffer for 5 minutes, washed with Wash Buffer (BRB-80 plus 20 μM paclitaxel), and blocked for 5 minutes with BRB-80 plus 5% w/v Pluronic F-127, followed by a second wash with Wash Buffer. Polymerized, rhodamine-labeled MTs were diluted 1:400 in BRB-80 plus 20 μM paclitaxel, flowed into the motility chamber for 10 minutes, and followed by a final wash with Wash Buffer. Kinesin-liposome complexes were diluted 1:20 into Motility Buffer (Buffer 1 with 2 mM MgATP, 20 μM paclitaxel, 0.5 mg/ml BSA, 0.5% w/v Pluronic F-127, 5 mM creatine phosphate, 0.4 mg/ml creatine phosphokinase, 10 mM DTT, 3.5 mg/ml glucose, 40 μg/ml glucose oxidase, 27 μg/ml catalase) and then perfused into the motility chamber. Still images of MTs and videos of liposome or Qdot motility were acquired using a custom-built TIRF microscope (22) with 532-nm excitation for MTs and 639-nm excitation for Qdots or liposomes. Motility videos were acquired at 10 frames per second.

#### Data analysis

To analyze movies of cargo motility on surface-bound microtubules, a line was drawn along a microtubule in the still MT image and duplicated to the cargo movie. Kymographs were generated (Multi Kymograph, ImageJ) where the x-axis is distance and y-axis is time so that moving cargoes appear as slanted lines in kymographs (Fig. 2A). To measure cargo velocity, a bounding box image was drawn to encompass the kymographic trajectory. Velocity (i.e. trajectory slope) was calculated as the box width (run length) divided by the box height (duration). For an event to be included in the run length analysis, it had to begin during the acquisition of the movie. Events which ended when the cargo reached the MT end were denoted as such. The “survival” package (63, 64) for R was used to compute the Kaplan-Meier estimate of the median run length for each experimental condition, accounting for termination events due to being track-limited.

### Liposome-kinesin-1 complex transport through 2D microtubule intersections

For 2D MT intersection assays, crossflow motility chambers were created by affixing a 22×22 mm no. 1 cover slip to a silanized 48×60 mm no. 1 cover glass with UV-curable adhesive and 125 μm Mylar shims. In this case, square shims were cut and the cover slip was mounted such that there were two perpendicular flow channels between the shims placed at each corner of the cover slip. The crossflow chamber was perfused with 0.8% w/v anti-tubulin antibody in BRB-80, both channels were washed with Wash Buffer and blocked with 5% w/v Pluronic F-127 in BRB-80. Following a second wash in each channel, dimly-fluorescent MTs diluted 1:400 in BRB-80 plus 20 μM paclitaxel was perfused into one channel, allowed to bind for 5 minutes, followed by a second perfusion of dim MTs into the same channel, followed by a wash with Wash Buffer. By being the first MTs to be infused into the chamber, dimly-fluorescent MTs would become the bottom MT of an intersection. Then, brightly-fluorescent MTs were diluted 1:400 in BRB-80 plus 20 μM paclitaxel and perfused into the perpendicular channel, allowed to bind for 5 minutes, followed by a second perfusion of the bright MTs into the same channel and a wash with Wash Buffer. Both channels were washed one final time with Wash Buffer, followed by a perfusion of kinesin-liposome complex diluted 1:20 in Motility Buffer. Still images of MTs and videos of liposomes were acquired as described above.

#### Data analysis

To analyze liposome motility and directional outcomes in 2D MT intersections, kymographs were used. In this case, both MTs defining the intersection were traced with segmented lines, and a kymograph of liposome transport corresponding to each MT was generated. The kymograph corresponding to the bottom, dimly-fluorescent MT was colored red, while the kymograph corresponding to the top, brightly-fluorescent MT was colored green. The two kymographs were then aligned to the intersection point and overlaid. While directional outcomes could be ascertained by eye in the liposome movies, pause lifetime in the intersection was measured by looking for and quantifying a vertical line (i.e. time axis) in the kymograph at the intersection point (see Fig. S7A for example from a 3D intersection).

### Optical trap ramp force assay

Ramp force measurements of kinesin-lipid-coated beads were made using a commercially available optical trapping system (Lumicks C-Trap System). 500 nm diameter lipid-coated beads were generated as reported previously (22), modified from (65, 66). Briefly, liposomes were prepared as described above, then diluted 3x into Buffer A (10 mM HEPES pH = 7.2 and 150 mM NaCl). Liposomes were sonicated (Fisher Scientific Sonic Dismembrator 550) with 0.5s alternating on/off pulses for a total elapsed time of 20 min (10 min total sonication) in an ice water bath, then were centrifuged at 5,000 x g for 10 min, and the supernatant saved in a fresh tube. 50 μl of 500 nm diameter silica beads (Duke Standard) was rinsed 3x with methanol and dried under vacuum prior to resuspension in 200 μl of Buffer A. Liposomes were incubated at 60°C for 2 min and mixed rapidly with the resuspended silica beads and immediately vortexed. Following vortexing, the lipid-bead mixture was shaken at low speed on a vortex mixer for 1 hour at room temperature to promote adsorption of the lipid onto the bead surface. Lipid-coated beads were then centrifuged at 5,000 x g for 3 min and resuspended in 200 μl of PBS (pH = 7.4) 3 times prior to final resuspension in 200 μl of PBS (pH = 7.4). Kinesin-543 was then affixed to the lipid-coated beads by incubation for 1 hour on ice at two different incubation ratios. For multi-motor ramp force assays, lipid-coated beads were incubated with a 20-fold molar excess of kinesin-543 to achieve the same density of motors as on liposomes with a 10-fold molar excess of kinesin-543. For single molecule ramp force assays, lipid-coated beads were incubated with limiting kinesin-543 such that ∼1 in every 10 lipid-coated beads screened would interact with the microtubule, generating force.

Ramp force experiments were performed in a custom flow cell generated by affixing a 22×22 mm no. 1 cover slip to a nitrocellulose-coated 24×60 mm no. 1 cover slip using UV-curable adhesive and 125 μm Mylar shims. Flow cells were first coated with 0.8% w/v anti-tubulin antibody in BRB-80 and allowed to incubate for 5 minutes prior to a wash with Wash Buffer. Then, the chamber was blocked with 1 mg/ml BSA in Buffer 1 for 5 minutes prior to a second wash with Wash Buffer. Rhodamine MTs were diluted 1:400 in BRB-80 plus 20 μM paclitaxel, perfused into the motility chamber, allowed to bind for 5 minutes, followed by a second perfusion and second 5-minute binding period, and a final wash with Wash Buffer. This strategy aligned the MTs to the direction of flow. Finally, the chamber was perfused with Motility Buffer, and 2 μl of 1:10 diluted lipid-coated beads in Motility Buffer were added. Individual beads were trapped, and a power spectrum was collected using the BlueLake software (Lumicks) to calibrate the trap stiffness for each bead. Trap stiffness ranged from 0.04 to 0.06 pN/nm. Beads were manipulated to be in close proximity to a rhodamine MT, and recordings of force ramp events due to kinesin-driven motion of the bead towards the MT plus end were collected. In multi-motor conditions, 213 events representing 8 beads from 3 independent bead preparations were collected, while for single motor conditions 34 events representing 7 beads from 3 independent bead preparations were collected.

#### Data Analysis

Force traces generated by kinesin-lipid-coated beads against the resistive load of the optical trap were analyzed to determine the maximum force attained during a ramp force event and its lifetime. Using a custom-written R script, median force was calculated using a rolling time-window (12.5 ms). The start of an event was identified when the median force reached 1 pN above baseline with the event termination (i.e., bead detachment from MT) identified by a rapid, single time step return to baseline, confirmed by a local maximum (≥2 pN) in a windowed (50 ms) rolling difference in force between time points. The detachment force attained during event was the median force over the 50 ms window prior to detachment, with a detachment force ≥ 2 pN considered above the noise. The event lifetime was the time elapsed between the start and end of the event. See Fig. 3B for example traces. Gaussian distributions were fit to the maximum force data using the fitdistr function of the MASS package (67) in R. Log-likelihood ratio testing in R (67) was used to determine if additional fit parameters were statistically justifiable.

### 3D microtubule tightrope and intersection assays

Crossflow motility chambers were created by affixing a 22×22 mm no. 1 cover slip to a plasma-cleaned 48×60 mm no. 1 cover glass using UV-curable adhesive and 125 μm Mylar shims. To suspend MTs off the motility chamber surface, silica microspheres with diameters ranging from 1.0 μm to 3.0 μm were used as pedestals. To promote MT binding to the silica beads, beads were coated with poly-L-lysine by pelleting gently (3,000 x g for 2 minutes) and resuspending in a solution of 400 μg/ml poly-L-lysine in 0.5 M TRIS Buffer (pH = 8.0). The beads were then incubated overnight under mild agitation, washed 3x (3,000 x g for 2 minutes) and resuspended in 1 M TRIS Buffer (pH = 8.0) and diluted to a final concentration of ∼1% solids. Beads were then perfused into the crossflow motility chamber and washed 2x each flow lane with 1M TRIS (pH = 8.0).

To suspend MTs between pedestal beads on the slide surface, a sequential flow approach was performed. First, both channels of the crossflow chamber were blocked with 1 mg/ml BSA in Buffer 1 for 2 minutes. Alexa-647 MTs were diluted 1:100 in BRB-80 buffer with 20 μM paclitaxel. Diluted MTs were flowed through the first channel and allowed to incubate for 2 minutes, followed by a second flow of diluted MTs, 2 minutes further incubation and a wash with BRB-80 plus 20 μM paclitaxel. This procedure was then repeated in the perpendicular channel, followed by a final wash with BRB-80 with 20 μM paclitaxel into both channels. 3D STORM imaging of the suspended MTs was performed in Buffer 1 with 50 mM beta-mercaptoethanol (βME), 20 mM Cysteamine (MEA) and 20 μM paclitaxel. Prior to performing motility experiments, the flow cell was washed twice with Motility Buffer. Kinesin-liposome complexes were diluted 1:100 in Motility Buffer and perfused into the flow cell prior to imaging.

3D motility experiments were imaged using a Nikon N-STORM microscope system. The system is equipped with a cylindrical lens and piezoelectric stage, which allowed for the calibration of the Z-axis (69, 70). To generate a STORM image of Alexa-647 MTs, 405 and 647 nm laser excitation was used, while 561 nm laser excitation was used to image DiI-labelled liposomes. To create the final reconstruction of the suspended MTs, a stack of 20,000 images was acquired at a framerate of ∼60 Hz. Liposome motility was imaged at a framerate of 10 Hz. Because the pedestal beads were visible in both the MT and liposome channels, they served as fiducial markers to ensure that the liposome image was in registry with the MT image in the XY plane. The built-in Nikon perfect focus system ensured that all images were in registry on the Z-axis.

#### Data analysis

Single-particle localizations for both MT STORM and liposome images were performed using the DoM (Detection of Molecules) plugin for ImageJ (71). The Z-calibration curve from the Nikon software was duplicated into DoM. The cross-correlation based drift correction tool built into DoM was used to remove the XY-drift from both liposome and MT STORM images. To link liposome localizations into a track defining each run, the “Link Particles to Tracks” function in DoM was used. To analyze liposome tracks along MT tightropes for helical trajectories, liposome tracks were reoriented such that the event ran along the X-axis in the plane Z=0. The event coordinates were then projected onto the X-Y and X-Z planes and fit with a sinusoid (Fig. S6A) and the sinusoids were combined to yield a 3D helix (Fig. 5A).

For each 3D MT intersection, an ROI (Region of Interest) was selected which contained the liposome track and the MTs which defined the intersection. The STORM localizations of the intersecting MTs were transferred to a custom R script which was used to fit a 3D line segment to the point cloud defining each MT (23). Then, a second R script overlaid the liposome trajectory onto the line segments defining the 3D MT intersection and characterized the motion of the liposome through the intersection and the Z-axis gap (*d*) between the intersecting MTs at the point of intersection. The angle between the two crossing MTs projected onto the x-y plane was calculated, and only intersections with angles between 60° and 120° were included. The liposome approach angle, α, was determined by averaging the Z position of the liposome center over 10 frames prior to it reaching the crossing MT. From these data an azimuthal angle of the liposome center relative to a perpendicular line drawn between the intersecting MT centers (0° axis) was calculated (Fig. 6A). To determine if liposomes paused in 3D intersections and measure the duration of such pauses, kymographs were used. We generated two kymographs, one for each of the intersecting MTs. The two kymographs were colored green and red, aligned such that the intersection point was in the same location for both kymographs, and overlaid. With position on the horizontal axis and time on the vertical axis, a pause was indicated by a vertical line in the kymographs at the intersection point. The duration of the pause was determined by measuring the vertical line length (i.e., length of time) in the kymograph at the intersection point (Fig. S7A). Only pauses of at least 3 frames (0.3 s) were included in the analysis as shorter pauses could not be visually identified with confidence.

## Acknowledgements

We would like to thank Shane R. Nelson for advice and support developing analysis software, M. Yusuf Ali for experimental advice, Guy Kennedy for support and training in TIRF microscopy, Andrew Lombardo for experimental and analysis advice, Patricia M. Fagnant for protein expression and purification, Douglas Taatjes and Nicole Bouffard of the UVM Microscopy Imaging Center (RRID# SCR_018821) for support and training in 3D NSTORM imaging, and current and former members of the Warshaw and Trybus labs for valuable input, support, and discussions. We acknowledge the contributions of those who have helped to create, distribute, and maintain the open-source software used in this study, specifically r-project.org, and ImageJ.nih.gov. This research was funded by NIH Grant T32HL076122 (to B.M.B.), NIH Grant F32GM140618 (to B.M.B.), NIH Grant R35GM141743 (to D.M.W.), NIH Grant R35GM136288 (to K.M.T.), and NIH Grant S10OD026884 (to D.M.W.), and a generous gift from Arnold and Mariel Goran to D.M.W..

## Notes

### Competing Interest Statement

The authors have declared no competing interest.

