## Supplementary Materials for "Spatial Relationships Matter: Kinesin-1 Molecular Motors Transport Liposome Cargo Through 3D Microtubule Intersections *In Vitro*"

**Supplementary Movies**

**Movie S1.** Liposome with ~10 bound kinesin-1 proceeds straight through 3D MT intersection. Liposome motility movie (yellow channel) is overlaid with 3D STORM reconstruction of suspended MTs (3D STORM Z-axis color scale shown in Fig. 1B of main text). MT-MT intersection gap is 250 nm. Movie playback 1X real time. Scale 500 nm.

**Movie S2.** Liposome with ~10 bound kinesin-1 turns in 3D MT intersection. Liposome motility movie (yellow channel) is overlaid with 3D STORM reconstruction of suspended MTs (3D STORM Z-axis color scale shown in Fig. 1B of main text). MT-MT intersection gap is 130 nm. Movie playback 1X real time. Scale 500 nm.

**Movie S3.** Liposomes with ~10 bound kinesin-1 turn and proceed straight equally in 2D MT intersections when approaching from the bottom MT. MTs (green) are made with dim and bright fluorescence labeling. Dim MTs are introduced into the chamber first and are thus the bottom MT in intersections, followed by introducing bright MTs, which become the top MT in the intersection. Two liposomes (red) approach the bright, top MT from the dim, bottom MT. The first turns at the intersection, while the second proceeds straight. Movie playback 2X real time. Scale 500 nm.

For **Movies S4** and **S5** depicting simulations of 500 nm lipid-coated beads with 20 kinesin-1 motors stepping against the resistive load of an optical trap. Lower movie: MT (green rod), trapped bead is translucent orange with a blue equatorial stripe to indicate orientation, kinesin-1 motors on the liposome surface not bound to a MT are purple spheres, and MT-bound kinesin-1 motors are represented by its tail (purple rod) connected to one head/neck in red and the other head/neck in yellow. The spring (red line) connecting the kinesin tail to the liposome represents the overall tail elasticity (see Supplementary Information). Time shown is elapsed time from when the first kinesin-1 motor attaches to the MT with the bead at trap center (i.e. no hindering load). N is the number of kinesin-1 motors bound to the MT at every point in time. F is the resistive trap force applied to the bead as the kinesin-1 motors attempt to transport the bead away from trap center. Upper movie: A close-up view of the motors at the front of the liposome with each ab-tubulin dimer (blue and white hemispheres) depicted in the MT. Playback speed 1/6$^{th}$ real time.

**Movie S4:** Simulation of kinesin-1-transported lipid-coated bead in the optical trap. Number of attached motors varies during simulation, reaching a maximum force of ~18 pN. At this point 3 kinesin-1 motors share the load. When the resisting load exceeds that generated by the attached motors, which occurs when one or more motors detach, the remaining motors slowly backstep, causing force to drop, until 3 motors again share the force and approach final stall force. Notice that motors can attach to different protofilaments.

**Movie S5:** Simulation of kinesin-1-transported lipid-coated bead in the optical trap. For the entire simulation only a single kinesin-1 motor steps against the resistive force of the optical trap, which reaches a maximum of ~7 pN. Such single motor transport events were the predominant scenario for the simulations.

For **Movies S6-S10** depicting simulations of 350 nm liposomes with 10 kinesin-1 motors encountering a MT-MT intersection with MTs as green rods, liposome is translucent orange with a blue equatorial stripe to indicate orientation, kinesin-1 motors on the liposome surface not bound to a MT are purple spheres, and MT-bound kinesin-1 motors are represented by its tail (purple rod) connected to one head/neck in red and the other head/neck in yellow. The spring (red line) connecting the kinesin tail to the liposome represents the overall tail elasticity (see Supplementary Information). Time shown is elapsed time since simulation initiation, with movies starting once the system has reached steady-state and prior to encountering the intersection, ~t=1.4s. Kinesin-1 motors start attached to a MT, moving toward its plus-end (to the right) with the crossing MT plus-end pointing away from the viewer. N is the number of kinesin-1 motors bound to the MTs at every point in time. Playback speed 1/6$^{th}$ real time.

**Movie S6:** Turn Directional Outcome in MT Intersection. When the crossing MT presents a significant barrier to transport, liposomes may turn at an intersection when the kinesin-1 motors on the crossing MT "win" the tug of war. This simulation shows an intersection gap (*d*) of 75 nm and a small approach angle. Like **Movie S8**, kinesin-1 motors attach to the crossing MT and initiate a tug of war, but "win" the tug of war when motors on the starting MT detach, allowing motors on the crossing MT to continue transporting the liposome away from the intersection, yielding a turn directional outcome. We observe this mechanism both for small and large microtubule separations (see **Movie S7**).

**Movie S7:** Turn Directional Outcome in MT Intersection. When the crossing MT does not present a significant barrier to transport, liposomes may turn at an intersection when the kinesin-1 motors on the crossing MT "win" the tug of war. This simulation shows an intersection gap (*d*) of 225 nm and a small approach angle. Like **Movie S9**, kinesin-1 motors attach to the crossing MT and initiate a tug of war, but "win" the tug of war when motors on the starting MT detach, allowing the motors on the crossing MT to continue transporting the liposome away from the intersection, yielding a turn directional outcome. We observe this mechanism both for small and large microtubule separations (see **Movie S6**).

**Movie S8:** Straight Directional Outcome in MT Intersection. When the crossing MT presents a significant barrier to transport, liposomes may proceed straight through an intersection when the liposome is pulled under the intersection during a tug of war. This simulation shows an intersection gap (*d*) of 75 nm and a small approach angle. Kinesin-1 motors attach to the crossing MT and initiate a tug of war. Eventually the liposome is pulled down through the intersection. The motors on the crossing MT "lose" the tug of war and detach, allowing motors on the starting MT to continue transporting the liposome past the intersection, yielding a straight directional outcome. We observe this mechanism both for small and large MT gaps (see **Movie S9**).

**Movie S9** Straight Directional Outcome in MT Intersection. When the crossing MT does not present a significant barrier to transport, liposomes may proceed straight through an intersection when the liposome is pulled under the intersection during a tug of war. This simulation shows an intersection gap (*d*) of 225 nm and a small approach angle. Kinesin-1 motors attach to the crossing MT and initiate a tug of war, eventually the liposome is pulled down through the intersection. The motors on the crossing MT "lose" the tug of war and detach, allowing motors on the starting MT to continue transporting the liposome past the intersection, yielding a straight directional outcome. We observe this mechanism both for small and large microtubule separations (see **Movie S8**).

**Movie S10:** Straight Directional Outcome in MT Intersection. When the crossing MT does not present a significant barrier to transport, liposomes may proceed straight through an intersection when the liposome diffuses under the intersection. This simulation shows an intersection gap (*d*) of 225 nm and a small approach angle. Kinesin-1 motors never attach to the crossing MT, allowing the liposome to move under the crossing MT by diffusion and to continue straight past the intersection, yielding a straight directional outcome. We observe this mechanism only for large microtubule separations.

**Supplementary Figures.**

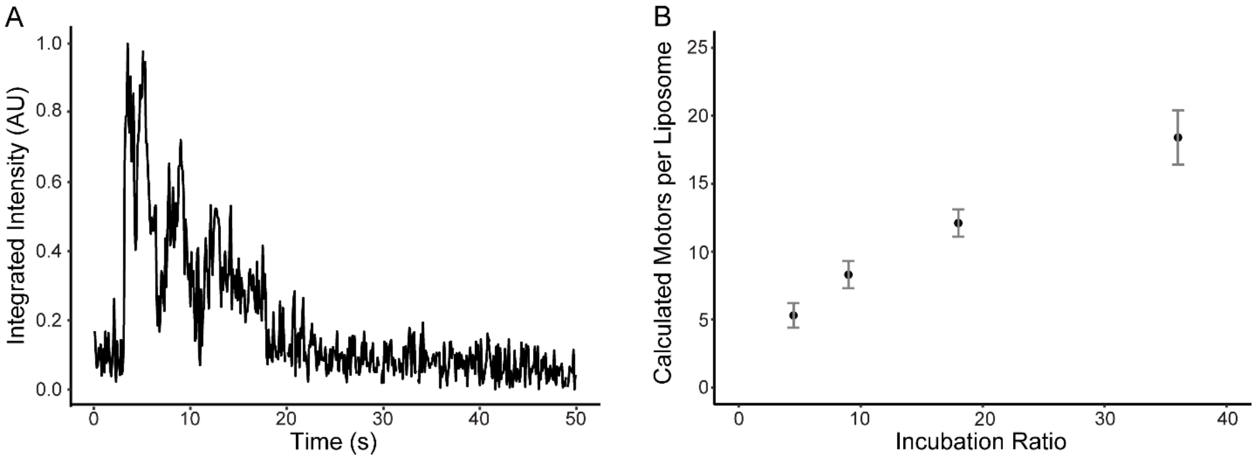


Figure S1. *Photobleaching Analysis to Estimate Kinesin Number per Liposome.* (A) Sample fluorescence decay curve of multiple YFP-kinesin attached to a liposome. (B) Plot of measured number of motors per liposome versus the incubation ratio (kinesin per liposome).

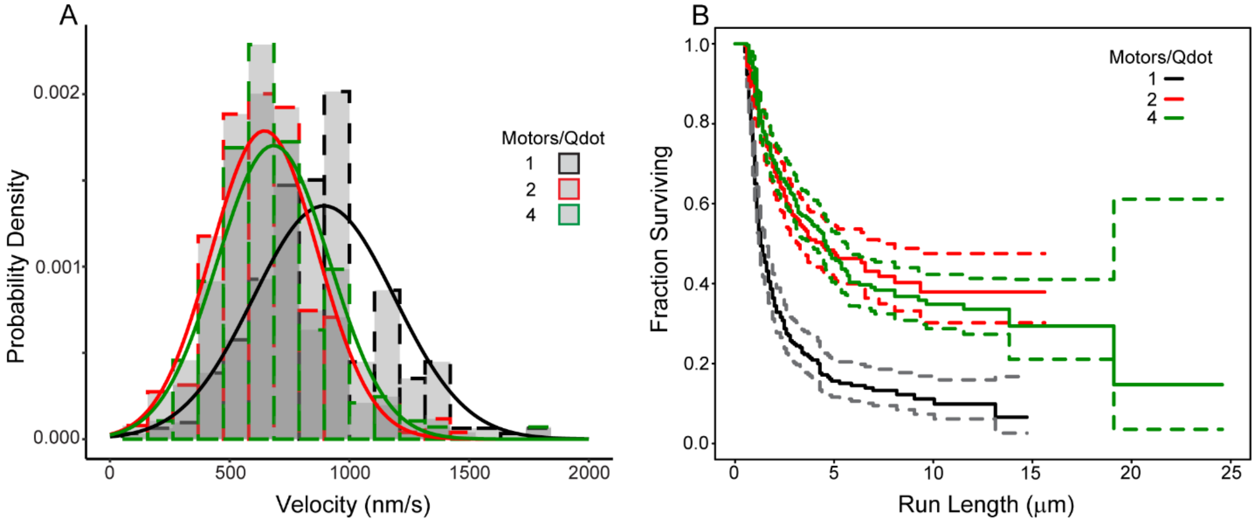


Figure S2. *Velocity and Run Length of Kinesin-Qdot Complexes Depend on Motor to Qdot Incubation Ratio.* (A) Histogram and Gaussian fit (solid curves) to velocity distribution for various motor to Qdot incubation ratios: 1:1 (890±20 nm/s, mean±SEM), 2:1 (650±10 nm/s, mean±SEM), and 4:1 (680±10 nm/s, mean±SEM). (B) Survival curve (solid curve) and 95% CI (dashed curves) of run length for various motor to Qdot incubation ratios: 1:1 (1.3 μm median), 2:1 (2.8 μm, median), and 4:1 (3.5 μm, median).

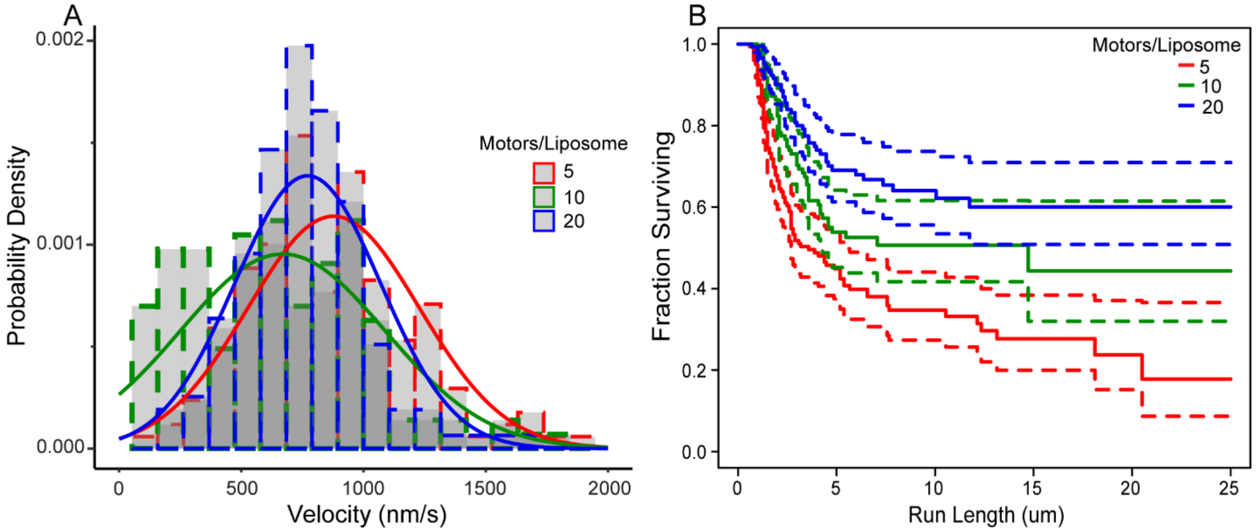


Figure S3. *Velocity and Run Length of Kinesin-Liposome Complexes Depend on Motor to Liposome Incubation Ratio.* (A) Histogram and Gaussian fit (solid curves) to velocity distribution for motor to liposome incubation ratios: 5:1 (800±20 nm/s, mean±SEM), 10:1 (670±40 nm/s, mean±SEM), and 20:1 (770±30 nm/s, mean±SEM). (B) Survival curve (solid curves) and 95% CIs (dashed curves) of run length for motor to liposome incubation ratios: 5:1 (3.6 μm median), 10:1 (14.7 μm, median) and 20:1(undefined, median).

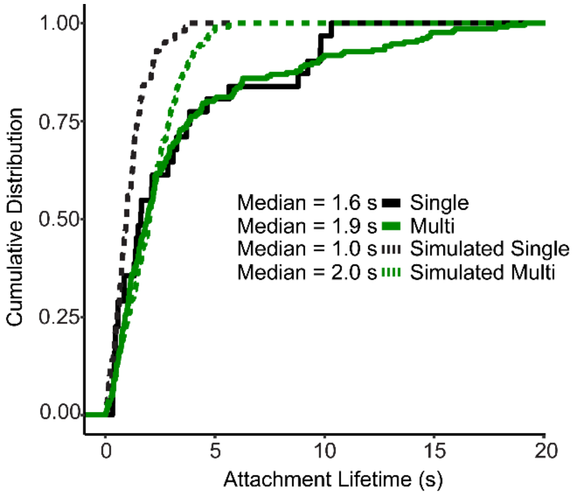


Figure S4. *Attachment Lifetimes Under Load Depend on the Kinesin-Bead Ratio.* Cumulative distribution plots of ramp force event lifetimes for single (black) and multiple (green) kinesins pulling against the trap resistive force. Lifetimes for simulated single (black dashed) and simulated multiple (green dashed) kinesins are also shown.

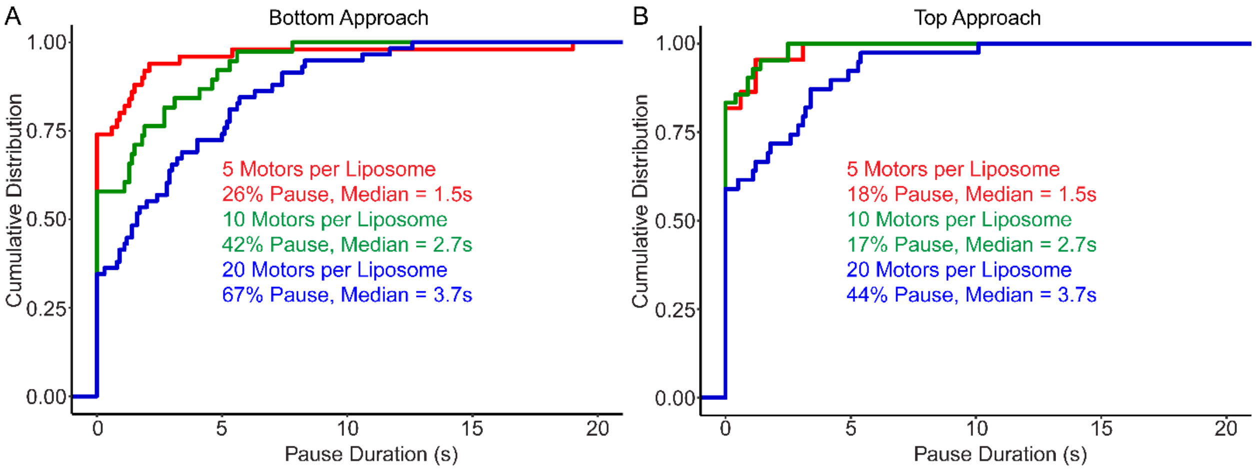


Figure S5. *Liposome Pause Frequency and Duration Depend on Ensemble Size and Approach MT in 2D MT Intersections.* (A) Cumulative distribution plot of observed pauses in 2D MT intersections where liposomes approach the intersection from the bottom MT. Different incubation ratios are shown by different colors. Text inset gives percentage of events where a pause was observed and the median duration of the observed pauses for each condition. (B) Cumulative distribution plot of observed pauses in 2D MT intersections where liposomes approach the intersection from the top MT. Different incubation ratios are shown by different colors. Text inset gives percentage of events where a pause was observed and the median duration of the observed pauses for each condition.

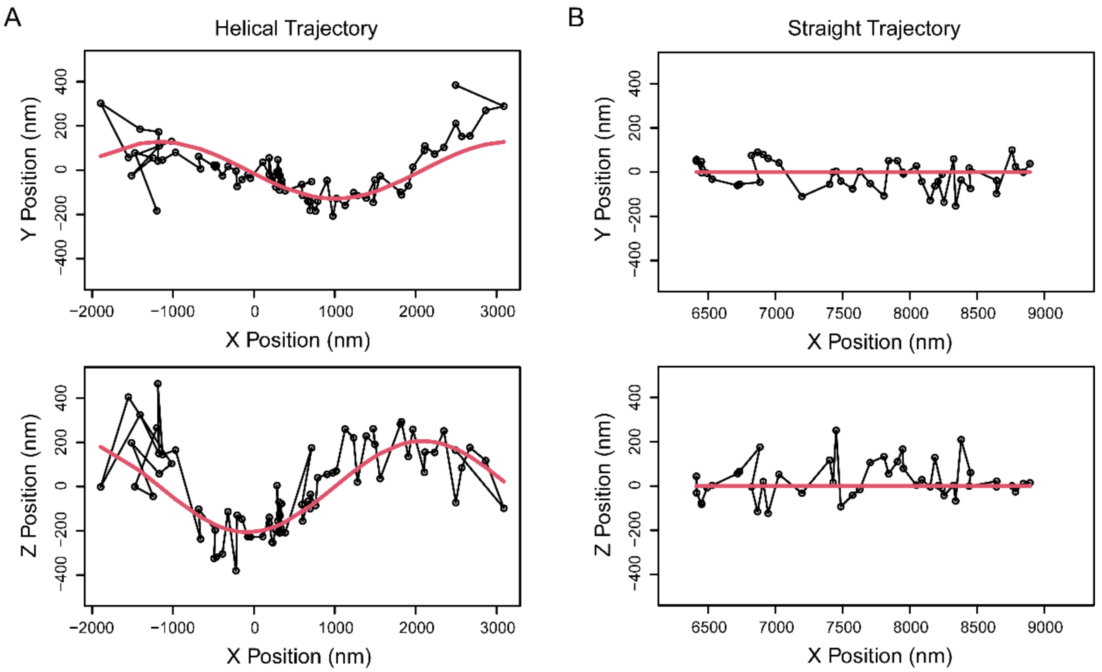


Figure S6. *X,Y and X,Z Projections of Liposome Trajectories Moving Along Suspended MTs Reveal Helical and Straight Motion.* (A) X,Y (top) and X,Z (bottom) projections of a helical liposome trajectory which has been transformed by mathematically centering the trajectory along the X-axis. Black dots represent the transformed liposome center localizations. Red curve represents the X,Y or X,Z component of a 3D sinusoid fit to the trajectory. (B) X,Y (top) and X,Z (bottom) projections of a straight liposome trajectory which has been transformed by mathematically centering the trajectory along the X-axis. Black dots represent the transformed liposome center localizations. Red line represents a line segment fit to the trajectory.

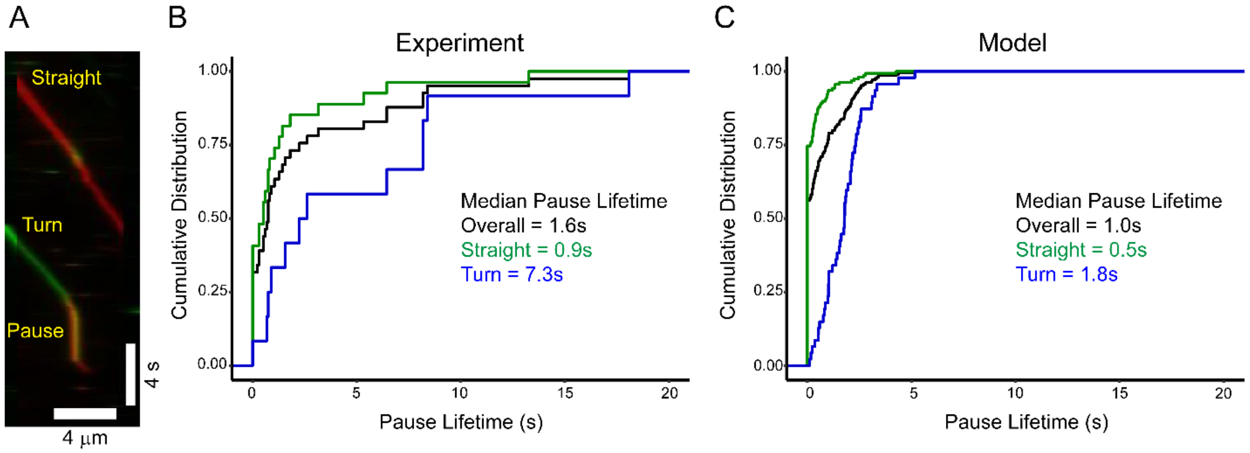


Figure S7. *Liposomes Pause in 3D MT Intersections More Frequently for Turn Outcomes than Straight Outcomes.* (A) 2-color kymograph of liposomes moving through a 3D MT intersection (see Methods for details). Time is represented along the vertical axis, and distance along the horizontal axis. Kymograph traces represent two different liposome trajectories separated in time encountering the same two intersecting MTs. Upper Trace: Liposome motion on the starting MT begins as a red trace and by going straight through the intersection remains red, except for the yellow spot where the liposome interacts with the crossing MT. Lower Trace: Liposome motion on the starting MT begins as a green trace but pauses at the intersection (yellow vertical trace) and then turns onto the intersecting MT, indicated by the red trace before terminating. (B) Cumulative distribution plot of measured pause lifetimes in the 3D MT intersection assay. Overall pauses regardless of directional outcome are black, pauses prior to going straight in green, and pauses prior to turning in blue. Median pause lifetimes exclude events that do not pause. (C) Cumulative distribution plot of modeled pause lifetimes in the 3D intersection simulation (overall (black), straight (green), and turn (blue). Median pause lifetimes exclude events that do not pause.

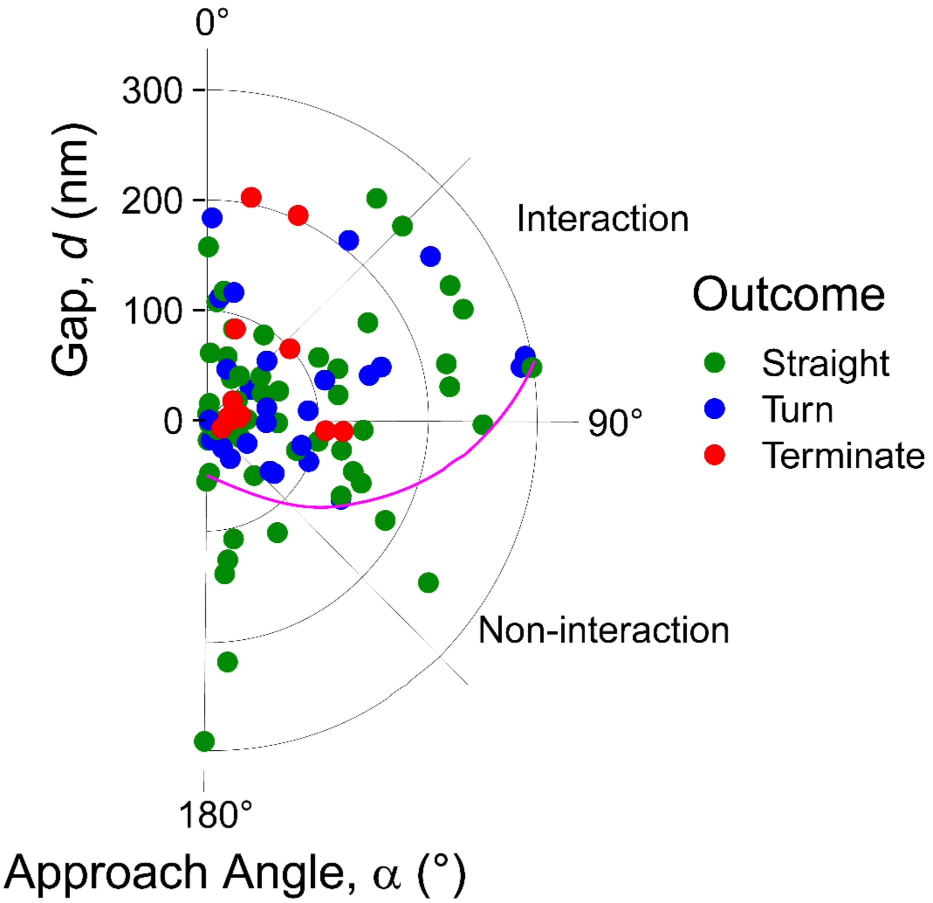


Figure S8. *Straight, Turn, and Terminate Outcomes Can Arise from Similar Interaction Geometries.* Polar scatter plot of straight (green), turn (blue), and terminate (red) outcomes as a function of approach angle ($\alpha$) and gap ($d$). Magenta curve defines border between zones where kinesin on the liposome surface can reach the intersecting MT and thus an interaction (above) or not and thus a non-interaction (below).

### Mathematical modeling supplement

Here we provide additional details of our mathematical modeling. Our approach is similar to previously published models [1–3], particularly recent 3-dimensional models with multiple motors [4–6]. There are, however, some important differences, mostly arising from modeling kinesin as a linear spring at all lengths, as opposed to a linear spring only in lengthening. For completeness, we include a full description of our modeling approach, highlighting differences between our model and previous models as they arise.

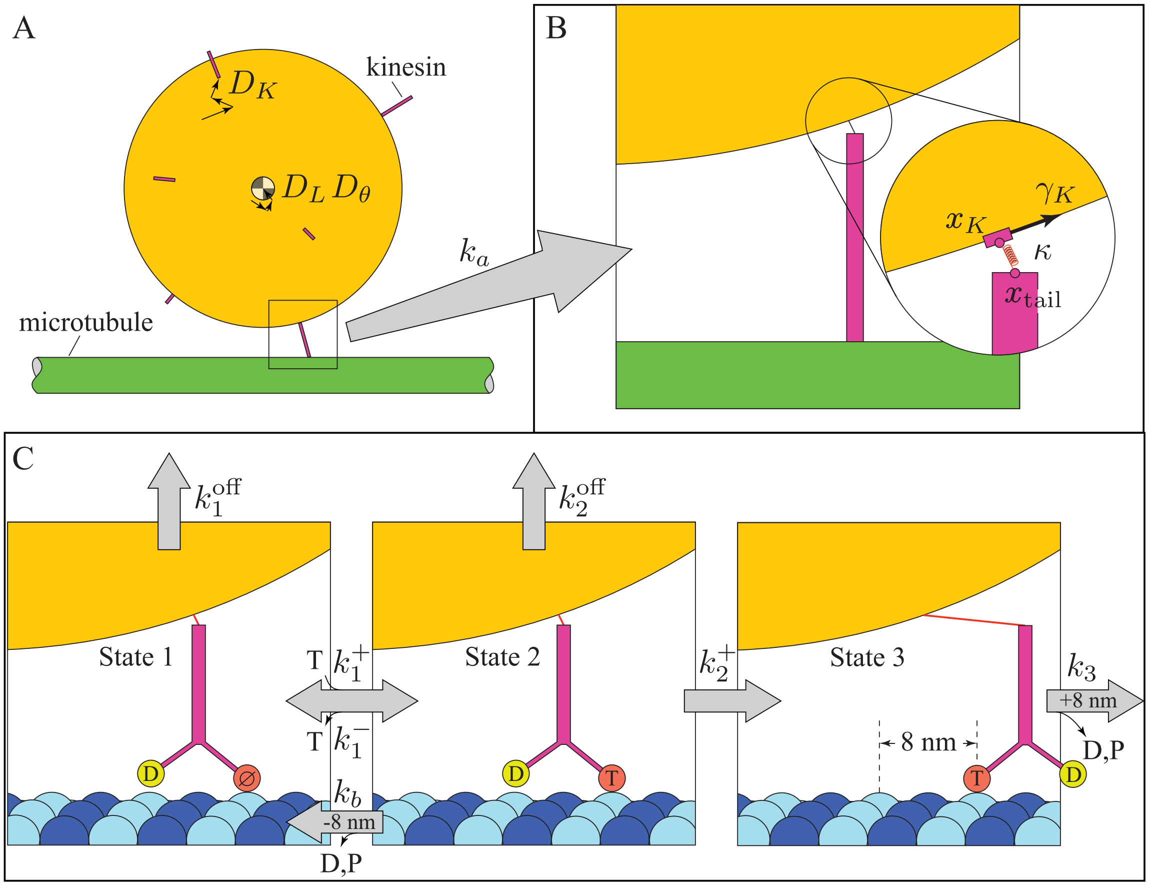


Figure S9: Mathematical model of microtubule-based transport of a fluid liposome by multiple kinesin. A. Kinesin (purple) diffuse with diffusion constant $D_K$ over the surface of the liposome (yellow). The liposome undergoes both linear and angular diffusion, with diffusion constants $D_L$ and $D_\theta$, respectively. When a kinesin molecule approaches the microtubule (green), it may attach. We have placed a box around one such potential interaction, which is pictured just prior to kinesin attachment. B. Once kinesin attaches to the microtubule, we model its attachment to the liposome by a linear spring, of stiffness $\kappa$, attached on one end to a ball joint on kinesin's tail (located at $\boldsymbol{x}_{\text{tail}}$) and on the other to a ball joint anchored in the liposome membrane (located at $\boldsymbol{x}_K$). This latter anchor can slide over the surface of the liposome with drag constant $\gamma_K$. C. Once kinesin attaches to the microtubule, we model its stepping and detachment according to a scheme based on single molecule experiments [12]. Both heads of the kinesin molecule are shown. To distinguish between the heads, one is shown in yellow and the other in red. Occupancy of the active site of each head is written inside. Note that the notation +8nm (i.e., forward step) returns kinesin to state 1, but with the heads switched (so the red one is where the yellow one is pictured and vice versa) and all positions moved forward 8nm. The notation −8nm (i.e., backward step) returns kinesin to state 1, but with the heads switched and all positions moved back 8nm. Abbreviations: ∅ = empty, T=ATP, D=ADP, P=inorganic phosphate.

### 1 Mechanics of kinesin, the liposome, and microtubules

Each kinesin molecule is modeled as a rigid rod 25nm in length (Fig. S9A). This length estimate comes from estimating kinesin's full length at $\sim$ 80nm, its heads at 10nm, and the consideration that our construct uses about a quarter of the tail (giving $0.25 \times (80 - 10)$nm + 10nm = 27.5nm $\approx$ 25nm) [7, 8]. It is attached to the liposome surface via a frictionless ball joint. There is an elastic element connecting the ball joint to kinesin's tail (the end of the rigid rod). This elastic element has a zero rest length, and stiffness $\kappa$ = 1pN/nm. This stiffness estimate comes from estimates of myosin V stiffness [9] that were successful in a similar model of multi-motor transport with myosin V [10, 11], and the expectation that this stiffness is approximately the stiffness of a coiled-coil tail, which would dominate the stiffness of our truncated construct. This spring applies a force, proportional to the distance to the ball joint, directed along its length (Fig. S9B). This elastic element models the bending and the extensibility of the kinesin molecule and the deformability of the liposome. The liposome is modeled as a rigid sphere. The ball joint of each kinesin molecule is anchored to the surface of the sphere, and slides over the surface with a viscous drag $\gamma_K$ (Fig. S9B). Collisions between the liposome and the microtubule(s) are perfectly elastic.

Each microtubule is modeled as a rigid cylinder, of radius 12.5nm. Kinesin molecules, when bound, extend orthogonally from the microtubule surface (Fig. S9B). Steric interactions between the kinesin molecules and the microtubules are neglected, as are steric interactions between kinesin molecules. In our microtubule intersection simulations, it is therefore possible for a kinesin molecule to "pass through" a crossing microtubule if the gap is too small ($d \approx$ 25nm), so the minimum gap in all simulations was set to 50nm.

### 2 Modeling fluctuations and dissipation

The dynamic motion of the liposome and of the kinesin molecules is modeled with an Euler-Maruyama scheme, as others have done previously [4–6]. The position of the liposome's center of mass, $\boldsymbol{x}_L(t)$, is advanced forward a time step $\Delta t$ with the equation

$$\boldsymbol{x}_L(t + \Delta t) = \boldsymbol{x}_L(t) + \frac{\Delta t}{\gamma_L} \sum_{i=1}^{N_B} \boldsymbol{F}_i + \boldsymbol{w}_f$$

where $N_B$ is the number of kinesin molecules bound to the microtubule(s), $\boldsymbol{F}_i$ is the force applied by the $i^{\text{th}}$ bound kinesin molecule on the liposome, $\gamma_L$ is the viscous drag of the liposome, and $\boldsymbol{w}_f$ is a term that captures position fluctuations from solvent collisions. In particular, $\boldsymbol{w}_f$ is a Gaussian random variable of mean 0 and standard deviation $\sqrt{2D_L\Delta t}$, where $D_L$ is the diffusion constant of the liposome.

Similarly, the orientation of the liposome, $\boldsymbol{\theta}_L(t)$, is advanced forward a time step $\Delta t$ with the equation

$$\boldsymbol{\theta}_L(t + \Delta t) = \boldsymbol{\theta}_L(t) + \frac{\Delta t}{\gamma_\theta} \sum_{i=1}^{N_B} \boldsymbol{M}_i + \boldsymbol{\phi}_f$$

where $N_B$ is the number of kinesin molecules bound to the microtubule(s), $\boldsymbol{M}_i$ is the moment applied by the $i^{\text{th}}$ bound kinesin molecule on the liposome, $\gamma_\theta$ is the angular viscous drag of the liposome, and $\boldsymbol{\phi}_f$ is a term that captures angular fluctuations from solvent collisions. In particular, $\boldsymbol{\phi}_f$ is a Gaussian random variable of mean 0 and standard deviation $\sqrt{2D_\theta\Delta t}$, where $D_\theta$ is the angular diffusion constant of the liposome.

Finally, the kinesin molecules diffuse over the surface of the liposome. Each obeys the equation

$$\boldsymbol{x}_K(t + \Delta t) = \boldsymbol{x}_K(t) + \frac{\Delta t}{\gamma_K} \left((\boldsymbol{F} \cdot \hat{\boldsymbol{e}}_{t1})\hat{\boldsymbol{e}}_{t1} + (\boldsymbol{F} \cdot \hat{\boldsymbol{e}}_{t2})\hat{\boldsymbol{e}}_{t2}\right) + w_1\hat{\boldsymbol{e}}_{t1} + w_2\hat{\boldsymbol{e}}_{t2}$$

where $\boldsymbol{F}$ is the force on the kinesin molecule (which is only non-zero for kinesin bound to microtubules), $\gamma_K$ is the viscous drag of kinesin moving across the liposome surface, $\hat{\boldsymbol{e}}_{t1}$ and $\hat{\boldsymbol{e}}_{t2}$ are any two orthogonal unit vectors originating from the point kinesin attaches to the liposome that are both tangent to the surface of the liposome, and $w_1$ and $w_2$ are Gaussian random variables of mean 0 and standard deviation $\sqrt{2D_K\Delta t}$, where $D_K$ is the diffusion constant of kinesin moving across the liposome surface. Note that, after we calculate $\boldsymbol{x}_K(t+\Delta t)$, we project back onto the surface of the sphere.

Each diffusion constant is related to the viscous drag constant through the relationship $D = k_BT/\gamma$, where $k_B$ is Boltzmann's constant and $T$ is temperature (assumed 25°C). For the liposome, we assume spherical Stokes drag so that $\gamma_L = 6\pi r_L\eta$, where $\eta = 0.001\cdot 10^{-6}\text{pN}\cdot\text{s/nm}$ is the dynamic viscosity of water at 25°C and $r_L$ is the liposome radius in nm. By the same assumption, the angular drag is $\gamma_\theta = 8\pi r_L^3\eta$. The diffusion constant of myosin V in the liposomes we used was $D_K = 0.92\cdot 10^6\text{nm}^2/\text{s}$ [13], which we assume is the same for kinesin, allowing us to calculate $\gamma_K = k_BT/D_K$.

### 3 Modeling chemical reactions

Each kinesin molecule follows a three state kinetic scheme [12], with a few modifications. The molecule starts in an apo state (state 1). It then binds ATP (state 2). Next, it steps forward (state 3), see Fig. S9C.

The ATP binding rate, $k_1^+$, governing the transition from state 1 to 2 is $k_1^+ = 3\mu\text{M}^{-1}\text{s}^{-1}$; the reverse rate is $k_1^- = 50\text{s}^{-1}$ [12]. When in state 1, kinesin may unbind from the microtubule in a force dependent process according to

$$k_1^{\text{off}} = \begin{cases} 0.1\text{s}^{-1} & : \text{ resistive force} \\ 0.9\text{s}^{-1} & : \text{ assistive force} \end{cases}$$

Like all force-dependent reactions in the scheme, this reaction depends on the component of force on kinesin that is directed along the microtubule, $F$.

The rate of forward stepping, governing the transition from state 2 to 3 is $k_2(F)$, and has the form $k_2^+(F) = k_{2,0}^+e^{-\delta F/k_BT}$. The rate of the release of hydrolysis products, governing the transition from state 3 back to state 1 (but moved forward 8nm) is $k_3$. We assume a back-stepping rate of $k_b = 5\text{s}^{-1}$. Because we allow back-steps, we independently estimated the parameters $k_{2,0}^+$, $\delta$, and $k_3$ by fitting measurements of force-velocity [12] with our model. In particular, we imported the measurements by clicking on each point in a PDF of the paper, and then fit the measurements using a non-linear optimization function in Matlab (fminsearch). The best fits were $k_{2,0}^+ = 2505\text{s}^{-1}$, $k_3 = 101.9\text{s}^{-1}$, and $\delta = 3.381\text{nm}$ (c.f., $k_{2,0}^+ = 2753\text{s}^{-1}$, $k_3 = 98.9\text{s}^{-1}$, and $\delta = 3.58\text{nm}$ [12], with no back-steps).

When in state 2, kinesin may unbind from the microtubule in a force dependent process according to

$$k_2^{\text{off}} = \begin{cases} k_{2r,0}^{\text{off}}e^{\delta_{2r}|F|/k_BT} & : \text{ resistive force} \\ k_{2a,0}^{\text{off}}e^{\delta_{2a}|F|/k_BT} & : \text{ assistive force} \end{cases}$$

We used $\delta_{2a} = 0.8\text{nm}$ [12]. We modified the value for resistive loads, using $\delta_{2r} = 0.7\text{nm}$ in our intersection simulations and $\delta_{2r} = 0.9\text{nm}$ in our laser trap simulations (the reported value is $\delta_{2r} = 1.1\text{nm}$ [12]). We justify these changes in a separate section, Section 10.3.

The values for $k_{2r,0}^{\text{off}} = 0.07\text{s}^{-1}$ and $k_{2a,0}^{\text{off}} = 0.13\text{s}^{-1}$ [12]. However, when we use these values we get single motor run lengths on the order of $5\mu\text{m}$, about three-fold higher than our measured single motor run length, $1.3\mu\text{m}$. We therefore increased these values three-fold ($k_{2r,0}^{\text{off}} = 0.21\text{s}^{-1}$, $k_{2a,0}^{\text{off}} = 0.39\text{s}^{-1}$), which brought the run lengths in our simulations in line with our measurements.

Finally, we allow the kinesin motors to (rarely) switch protofilaments. We added this to the model to capture observations of single kinesin molecules maneuvering around obstacles [16]. We anticipate that motors encountering a microtubule might switch protofilaments in a similar way, so we implemented it by allowing a protofilament switch once every 50 steps. This is done by selecting a random number from a uniform distribution at each kinesin step and, when the random number is less than 1/50, a sidestep is implemented to a neighboring protofilament, to the right or left with equal probability.

### 4 Attachment

The attachment of kinesin to microtubules is a chemical reaction, but we place it in a separate section to emphasize differences in our approach and some previous approaches [3–6]. In particular, we model the attachment rate of a kinesin molecule to a particular binding site on a microtubule with the equation

$$k_a = k_{a,0} \exp\left(-\frac{\kappa\left(\boldsymbol{x}_K - \boldsymbol{x}_{\mathrm{tail}}\right)^2}{2k_BT}\right)$$

where $\boldsymbol{x}_K$ is the position of the end of the kinesin molecule, $\boldsymbol{x}_{\mathrm{tail}}$ is the position of the tail of the kinesin molecule were it to bind to the binding site, and $\kappa$ is kinesin's assumed linear stiffness (Fig. S9B). The rate $k_{a,0}$ is the attachment rate if the binding site and end of the kinesin molecule are perfectly aligned. We assume $k_{a,0} = 150\mathrm{s}^{-1}$.

This differs from what previous authors have used, where binding rate is a constant if the point where a given kinesin molecule is attached to the liposome is within some radius of a binding site, and is zero otherwise. This approach gives each kinesin molecule a constant binding rate within a sphere centered at the liposome attachment point. By contrast, our approach gives each kinesin molecule a graded binding rate, with a maximum on a thin shell centered at the liposome attachment point, and then falling off as a Gaussian for both larger and smaller distances. This effectively reduces the overall attachment rate. Moreover, we expect it to introduce negative cooperativity, as kinesin binding tends to pull the liposome closer to the microtubule and makes it harder for unbound kinesin molecules to orient themselves in a position to maximize the attachment rate [10]. This contrasts with positive cooperativity predicted by models assuming a constant binding rate within a sphere, where kinesin binding tends to pull the liposome closer to the microtubule and makes it easier for unbound kinesin molecules to orient themselves in a position to maximize the attachment rate [5, 6].

In the model, we assume every microtubule has 13 protofilaments and a 3-start geometry. Each microtubule is assumed to have kinesin binding sites every 8nm along each of its protofilaments. These binding sites are 12.5nm from the center of the microtubule.

In principle, every kinesin molecule diffusing on the surface of the liposome could bind to each of the thousands of binding sites on each microtubule in our simulations. The vast majority of these binding reactions, however, would require the kinesin molecule to stretch a large distance and are therefore so unlikely as to be negligible. To improve the efficiency of our simulations, we therefore assume that a kinesin molecule will not bind to a binding site greater than 15nm away from its head, because the attachment rate for these reaction is less than $k_a < 2.37 \cdot 10^{-7}\mathrm{s}^{-1}$.

### 5 Numerically implementing the model

In the Euler-Maruyama scheme we use to approximate the stochastic dynamics of the liposome and kinesin motors, we must specify a time step, $\Delta t$. This value must be selected with care. In particular, it is critical that a typical diffusion step be small compared to the kinesin molecule, so that molecules will not take steps larger than the attachment zone. Additionally, we found that when our time step was too large, the simulations can become numerically unstable when multiple motors are attached and applying forces to each other. We therefore chose a very small time step, $\Delta t_1 = 5 \cdot 10^{-7}\mathrm{s}$.

With this time step, the expected change in liposome position each time step due to diffusion is 1.12nm and the expected position change due to diffusion for kinesin on the liposome surface is 0.96nm. The width of the attachment region is $\sqrt{k_BT/\kappa} = 2.03$nm, greater than these expected step sizes, so that there are two time steps on average within the attachment region.

A second potential problem is error introduced by round-off with a time step that is too small. That is, we simulate chemical reactions with a constant step method (see below). To do so, we estimate the probability of a given event happening in a time step and then compare that to a random number drawn from a uniform distribution between 0 and 1. If the probability is too small, then round-off error (where numbers below $10^{-14}$ are treated as zero) can skew simulation results. While our simulations would not likely be in this regime even if we used the time step of the Euler-Maruyama scheme, we used a larger time step for the chemical reactions $\Delta t_2 = 1 \cdot 10^{-5}$s – that is, we update the chemical reactions once every 20 Euler-Maruyama steps, which additionally reduces the computational cost of the simulations.

We simulate the chemical reactions with a constant step method, as opposed to more computationally efficient methods based on the Gillespie Algorithm, because we use a constant step in our Euler-Maruyama scheme. Additionally, it is the very large number of steps for the Euler-Maruyama scheme that limits our computation (e.g., we must take $10^7$ steps for a 5s simulation), so the computational savings from implementing the Gillespie algorithm would not have a large effect on total simulation run time.

The constant step method works as follows. Suppose there is a chemical reaction that occurs with rate constant $k$. In a given (small) time step $\Delta t$, we estimate the probability that the reaction occurs as $k\Delta t$. For every molecule that could potentially undergo the reaction, we draw a random number from a uniform distribution between 0 and 1 and, if the random number is less than the reaction probability $k\Delta t$, we implement the chemical reaction.

Because of the small time step in our simulations and the long simulation times, the output of our simulations would be unmanageable if we recorded, say, liposome position at each time point. For example, a 10 second simulation takes $2{\cdot}10^7$ time steps, so the position variable would be around 60Mb. Therefore, we define a time vector at the beginning of each simulation, typically with 4001 evenly-spaced points between 0 and 25s, and only record variables at those times. In this way, we keep all variables a manageable size.

### 6 Initial conditions

We chose to start all simulations with a single kinesin motor bound to the microtubule. All of the other kinesin molecules initially occupy the same space (N.B., there are no steric interactions between kinesin molecules in the simulation). To spread them across the surface of the liposome, we do not allow attachment to occur for the first 0.002s of each simulation. This allows each kinesin molecule to move an expected distance of 60nm from its initial position. Given that typical single motor runs are around 2s in duration, this small initial non-uniformity in the kinesin distribution and short period where motors may not attach is negligible.

### 7 Single microtubule simulations

We simulated runs of 350nm-diameter liposomes on a single microtubule with $N = 1$, $N = 5$, and $N = 10$ kinesin motors. All simulations started with a single kinesin motor bound, and proceeded until all kinesin molecules unbound or a maximum simulation time of 15s was reached.

For simulations with a single motor on the liposome, all simulations terminated prior to 15s. The median run length was 1.73$\mu$m ($n = 50$ simulations), though this includes simulations that detached prior to 0.3s (and so would not be counted as a run in our measurements). Excluding these simulations, the median run length was 1.91$\mu$m ($n = 46$ simulations). The median average speed (run length/attachment time) of these simulations was 776nm/s ($n = 46$ simulations).

For simulations with five motors on the liposome, five simulations were not complete by the 15s cutoff. Including these, the median run length was 3.36$\mu$m ($n = 50$ simulations). Excluding simulations that

Table S1: Model Parameters

| Param. | Description | Value/Equation | Reference/justification |
|---|---|---|---|
| $k_BT$ | Boltzmann's constant times temp. | $4.14\text{pN}\cdot\text{nm}$ | Calculated (25°C) |
| $\ell_K$ | Kinesin length | 25nm | [7,8] |
| $\kappa$ | Kinesin stiffness | 1pN/nm | [9] |
| $D_K$ | Diffusion of kinesin on the liposome | $0.92\cdot10^6\text{nm}^2/\text{s}$ | [13] |
| $\gamma_K$ | Drag of kinesin on the liposome | $4.5\cdot10^{-6}\text{pN}\cdot\text{s/nm}$ | $\gamma_K = k_BT/D_K$ |
| $r_L$ | Radius of liposome | 175nm* | see Methods |
| $\eta$ | Dynamic viscosity of water | $0.001\cdot10^{-6}\text{pN}\cdot\text{s/nm}^2$ | (25°C) |
| $\gamma_L$ | Drag on the liposome | $3.3\cdot10^{-6}\text{pN}\cdot\text{s/nm}$ | $\gamma_L = 6\pi r_L\eta$ |
| $D_L$ | Diffusion of liposome | $1.26\cdot10^6\text{nm}^2/\text{s}$ | $D_L = k_BT/\gamma_L$ |
| $\gamma_\theta$ | Angular drag on the liposome | $0.13\text{pN}\cdot\text{s}\cdot\text{nm}$ | $\gamma_\theta = 8\pi r_L^3\eta$ |
| $D_\theta$ | Angular diffusion of the liposome | $30.7\text{s}^{-1}$ | $D_\theta = k_BT/\gamma_\theta$ |
| $r_{MT}$ | Radius of microtubule | 12.5nm | |
| $k_1^+$ | ATP binding | $3\mu\text{M}^{-1}\text{s}^{-1}$ | [12] |
| $k_1^-$ | Reverse ATP binding | $50\text{s}^{-1}$ | [12] |
| $k_b$ | Back-stepping | $5\text{s}^{-1}$ | [14,15] |
| $k_2^+$ | Stepping | $k_{2,0}^+\exp\left(\frac{-\delta F}{k_BT}\right)$ | [12] |
| $k_{2,0}^+$ | Unloaded stepping | $2505\text{s}^{-1}$ | Fit to F-V from [12] |
| $\delta$ | Stepping force dependence | 3.381nm | Fit to F-V from [12] |
| $k_3$ | ADP/P$_\text{i}$ release | $101.9\text{s}^{-1}$ | Fit to F-V from [12] |
| $k_1^{\text{off}}$ | Unbinding, state 1 | $\begin{cases} 0.1\text{s}^{-1} & : \text{ resistive force} \\ 0.9\text{s}^{-1} & : \text{ assistive force} \end{cases}$ | [12] |
| $k_2^{\text{off}}$ | Unbinding, state 2 | $\begin{cases} k_{2r,0}^{\text{off}}\exp\left(\frac{\delta_{2r}\lvert F\rvert}{k_BT}\right) & : \text{ resistive force} \\ k_{2a,0}^{\text{off}}\exp\left(\frac{\delta_{2a}\lvert F\rvert}{k_BT}\right) & : \text{ assistive force} \end{cases}$ | [12] |
| $\delta_{2a}$ | Force dependence of unbinding, assistive | 0.8nm | [12] |
| $k_{2a,0}^{\text{off}}$ | Unloaded unbinding, assistive | $0.39\text{s}^{-1}$ | 3 times [12]† |
| $\delta_{2r}$ | Force dependence of unbinding, resistive | 0.7nm or 0.9nm | See section 10.3 |
| $k_{2r,0}^{\text{off}}$ | Unloaded unbinding, resistive | $0.21\text{s}^{-1}$ | 3 times [12]† |
| $k_a$ | Attachment rate | $k_{a,0}\exp\left(-\frac{\kappa(\boldsymbol{x}_K-\boldsymbol{x}_{\text{tail}})^2}{2k_BT}\right)$ | See section 4 |
| $k_{a,0}$ | Unloaded attachment | $150\text{s}^{-1}$ | See section 10.2 |
| $p_{switch}$ | Sidestep probability | 0.02 | [16] |

*We also used $r_L = 250\text{nm}$, adjusting other parameters $(\gamma_L, D_L, \gamma_\theta, D_\theta)$ affected by this change.
†Modified to explain single-motor binding duration and run length, see section 10.1.

detached prior to 0.3s, the median run length was 3.68$\mu$m ($n$ = 46 simulations). The median average speed (run length/attachment time) of these simulations was 776nm/s ($n$ = 46 simulations) – nearly identical to the $N = 1$ simulations.

For simulations with ten motors on the liposome, 29 simulations were not complete by the 15s cutoff. Including these, the median run length was 11.2$\mu$m ($n$ = 50 simulations). Excluding simulations that detached prior to 0.3s, the median run length was 11.3$\mu$m ($n$ = 47 simulations). The median average speed (run length/attachment time) of these simulations was 773nm/s ($n$ = 47 simulations).

#### 8 Laser trap simulations

To simulate our experiments in which kinesin motors transport a lipid-coated bead again the resistive force of the laser trap, we simulated runs of 500nm-diameter liposomes with $N = 20$ and with $N = 1$ kinesin motors on a single microtubule. All simulations started with a single kinesin motor bound, and proceeded until all kinesin molecules unbound or a maximum simulation time of 15s was reached. No simulations reached the maximum simulation time. We modeled the force of the laser by applying a force to the liposome's center of mass, directed along the microtubule toward the minus end, proportional to the distance traveled, $d_T$: $F = -k_{trap} d_T$. We assume a trap stiffness of $k_{trap} = 0.04$pN/nm.

#### 9 Microtubule intersection simulations

To simulate our experiments in which we assessed directional outcome at microtubule intersections, we simulated runs of 350nm-diameter liposomes with $N = 10$ kinesin motors. All simulations started with a single kinesin motor bound to one microtubule (the "starting" microtubule). The second microtubule (the "crossing" microtubule) was oriented perpendicular to the starting microtubule with its center a height $d$ above the center of the starting microtubule. The center of the crossing microtubule was positioned 1.6$\mu$m toward the plus end of the starting microtubule, to ensure that the system was in steady-state prior to the liposome entering the microtubule intersection. We oriented the crossing microtubule with its plus end to the left. In our simulations, there is nothing that breaks symmetry for the kinesin motors and the liposome (i.e., all chemical reactions and diffusion are equally likely to the left and right along the microtubule). The binding sites on the microtubule do have a handedness, but we anticipate that this has only a small effect so we only considered one orientation of the crossing microtubule. Simulations proceeded until 1) all kinesin molecules unbound, 2) a maximum simulation time of 15s was reached, 3) the liposome center of mass was 4 liposome diameters (1.4$\mu$m) past the intersection on either the starting or crossing microtubule. No simulations reached the maximum simulation time.

We simulated microtubule spacings of $d = 50, 75, 100, 125, 150, 175, 200, 225,$ and 250nm. For each spacing, we simulated the trajectories of 30 liposomes. Ten of these started with the attached kinesin directly on top of the starting microtubule on protofilament 1, giving an initial angle of 0. Note that this angle is not identical to the approach angle, since the simulated liposome started 1.6$\mu$m from the center of the crossing microtubule and the approach angle is assessed when the liposome is within 212.5nm of the center of the crossing microtubule, as described below. Ten of the simulations started with the attached kinesin on protofilaments 2 or 13, with equal probability, giving an initial angle of $\pm$ 25.7°. The final ten started with the attached kinesin on protofilaments 4 or 10, with equal probability, giving an initial angle of 77.1° or $-102.8$°. In this way, we sampled a range of approach angles and also varied the position of the kinesin motors on the microtubule.

For each of the 270 simulations, we determined the outcome as follows. If the liposome detached prior to encountering the intersection, it was not included in the analysis. This occurred for 60 simulations. The remaining 210 simulations were scored as a turn outcome if the liposome proceeded through the intersection on the crossing microtubule (47 simulations). They were scored as a terminate outcome if the liposome detached at the intersection (5 simulations). They were scored as a straight outcome if the liposome proceeded through the intersection on the starting microtubule (158 simulations).

To determine the approach angle, we assumed that the liposome reached the intersection when it proceeded to within the liposome radius (175nm) plus the motor length (25nm) of the surface of the crossing microtubule (i.e., within $r_L + \ell_K + r_{MT} = 175 + 25 + 12.5 = 212.5$nm of the center of the crossing microtubule). Placing a coordinate system with an origin located at the center of the starting microtubule with the $x$-axis pointing towards its plus end and the $z$-axis orthogonal to the crossing microtubule's center line, we then calculated the average height of the liposome, $\bar{z}_L$ over the previous 0.6s. The approach angle was $\alpha = \text{acos}(\bar{z}_L/r_{eff})$, where $r_{eff} = 182.5$nm. We determined the effective radius, $r_{eff}$ by trial and error, attempting to reproduce the actual approach angle (calculated from the simulated liposome's exact $x$, $y$ and $z$ values).

To determine pause duration at an intersection, we attempted to identify the beginning of the pause by looking at a plot of $x(t)$ and judging where it initially became constant. For straight outcomes, the end of the pause was determined by judging where $x(t)$ ceased being constant. For turn outcomes, we additionally plotted $y(t)$ and the end of the pause was determined by judging where $y(t)$ ceased being constant. The beginning of the pause was often difficult to identify with great confidence, while the end of the pause was clear because there is usually a jump in liposome position at the end of the tug of war. We therefore estimate an uncertainty of around 0.2s for each pause duration.

### 10 Parameter estimates

#### 10.1 Justification for changing $k^{\text{off}}_{2a,0}$ and $k^{\text{off}}_{2r,0}$

In our simulations of liposome transport along a single microtubule, we increased the values of $k^{\text{off}}_{2a,0} = 0.13\text{s}^{-1}$ and $k^{\text{off}}_{2r,0} = 0.07\text{s}^{-1}$ [12] three-fold. Without this change, the median run length for a liposome transported by a single kinesin was 6.09$\mu$m ($n = 100$ simulations). This run length is over 4-fold larger than what we measured with single, quantum dot-labeled kinesin (1.3$\mu$m). Our three-fold increase in these values reduces the median run length more than a factor of three, and brings it in line with our measurements.

#### 10.2 Justification for $k_{a,0}$

Kinesin's attachment rate to microtubules is determined by the parameter $k_{a,0}$, the attachment rate when the binding site and end of the kinesin molecule are perfectly aligned. In the single microtubule simulations, $k_{a,0}$ affects the run length of the simulations with 5 and 10 kinesin motors. To estimate $k_{a,0}$, we did a series of simulations of transport along a single microtubule with 10 kinesin motors (Fig. S10).

We performed 50 simulations with attachment rates of $k_{a,0} = 75, 150$, and $300\text{s}^{-1}$. Each simulation modeled transport of a liposome along a single, infinite microtubule, starting with a single kinesin bound, and ending when either 1) all kinesin motors detached from the microtubule, or 2) the simulation performed 30 million diffusion time steps (1.5 million chemical reaction time steps, or 15 seconds). To estimate $k_{a,0}$, we compared these simulation results to our experimental measurements. Since our measurements of liposomes with 10 kinesin motors often detached at the end of a microtubule, and because the distance to the end of the microtubule varied in our measurements, we had to estimate whether the simulated liposome would have reached the end of the microtubule. We did this by estimating the distribution of microtubule lengths in our measurements, picking lengths randomly from this distribution, and then comparing that length to our simulated run length.

To estimate the distribution of microtubule lengths in our measurements, we fit a function of the form

$$c(\ell) = \begin{cases} 1 - e^{-a(\ell-\ell_0)} & : \ \ell \geq \ell_0 \\ 0 & : \ \ell < \ell_0 \end{cases} \tag{1}$$

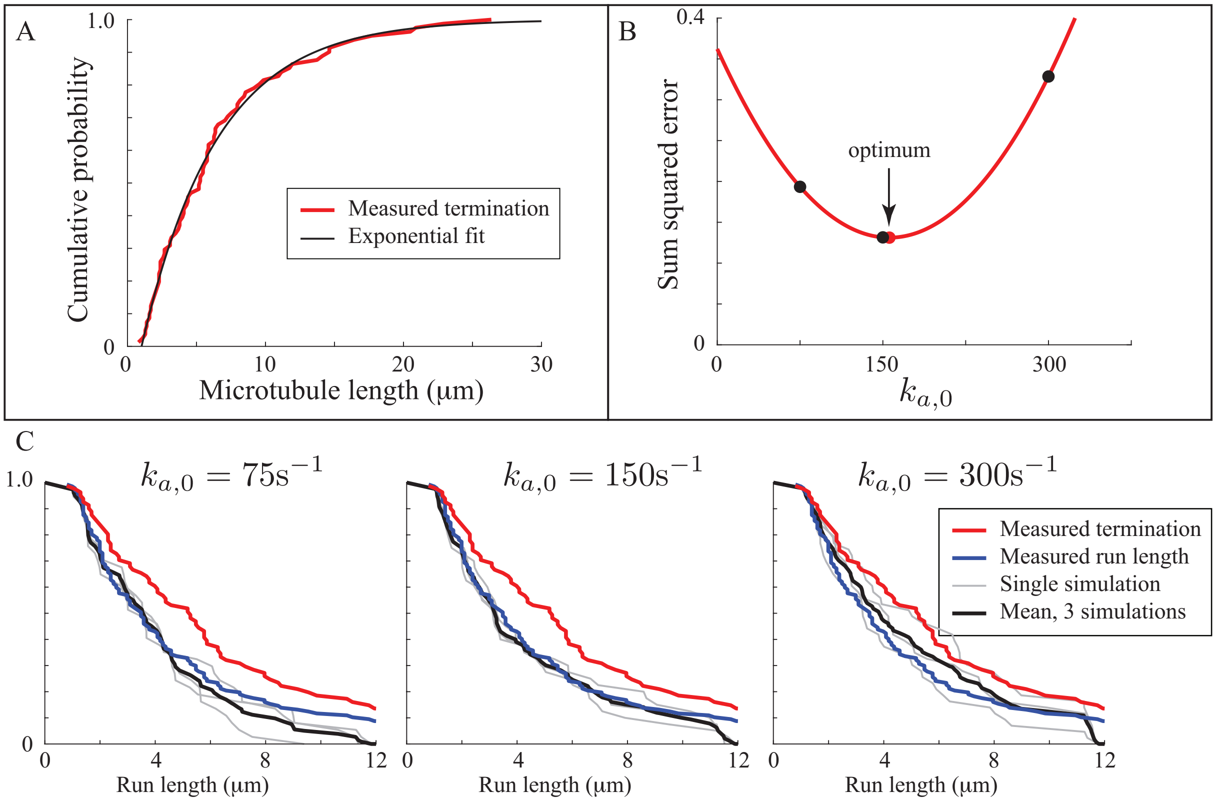


Figure S10: Measured run lengths of liposomes transported by 10 kinesin motors along a single microtubule predict an attachment rate of approximately $k_{a,0} = 150\text{s}^{-1}$. A. Estimating microtubule lengths by fitting an exponential curve, Eq. 1 (black), to our measurements that terminated at the end of a microtubule (red). The parameters of this best-fit are used to predict how far a simulated liposome may be transported by kinesin motors until it reaches the end of the microtubule. B. The sum of the squared error between simulated and measured run lengths (black dots) shows a minimum near $k_{a,0} = 150\text{s}^{-1}$. Constructing a quadratic curve (red) to pass through the three points allows us to estimate an optimal value of $k_{a,0} = 156\text{s}^{-1}$ (red dot). C. The curves that generated the points in panel B. Each gray line is the same set of 50 simulations on a different set of 50 microtubule lengths, the black line is the mean of these three. Note that when $k_{a,0}$ is too small (left curve), the simulations fall below the measurements, while when $k_{a,0}$ is too large (right curve), the simulations fall above the measurements. The red line is what run length would be if all liposomes traveled to the end of the microtubule.

to the experimentally measured cumulative probability distribution of run lengths of liposomes that detached at the end of the microtubule (Fig. S10A). Then, to pick a microtubule length for a given simulation, we used the best-fit values of the parameters $a$ and $\ell_0$ in the equation $\ell_i = -(1/a) \cdot \log(X_i) + \ell_0$ where $\ell_i$ is the length of the microtubule for the $i^{\text{th}}$ simulation, and $X_i$ is a random variable drawn from a uniform distribution between 0 and 1.

Once a microtubule length was determined for a simulation, we determined whether the liposome detached at the end of the microtubule. In particular, if the simulated run length was longer than the microtubule, then we assume detachment occurred at the end of the microtubule; otherwise, we assume detachment occurred at the end of the simulated run. In this way, we generated a set of simulated run lengths comparable to our measured run lengths. For each simulation, we generated microtubule lengths as described above three times. Calculating the mean squared difference between measurement and simulation for each of the three values of $k_{a,0}$, showed that $k_{a,0} = 150\text{s}^{-1}$ was the best-fit (Fig. S10B). Using the three points to define a quadratic curve, we estimate an optimal value of $k_{a,0} = 156\text{s}^{-1}$ (Fig. S10B), which is sufficiently close to $150\text{s}^{-1}$ that we used this latter value in our simulations.

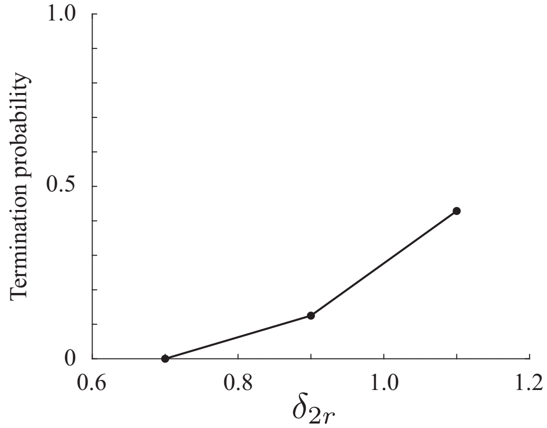


Figure S11: Reducing the force dependence of unbinding decreases termination at microtubule intersections. With a value of $\delta_{2r} = 1.1$nm [12], nearly half of simulations terminate at a separation of $z = 50$nm. Decreasing the load dependence to $\delta_{2r} = 0.7$nm eliminates these terminations, consistent with our experimental observations.

##### 10.3 Justification for reducing $\delta_{2r}$, force dependence of unbinding

When we used the value of $\delta_{2r} = 1.1$nm [12] in our simulations, liposomes frequently terminated their trajectory at microtubule intersections (Fig. S11). We did not observe this behavior in our measurements. In particular, we observed no termination outcomes for microtubule separations between 35 and 65nm, $n = 16$. We reasoned that the reported value force dependence [12] might reflect some of kinesin's sensitivity to vertical forces that are present in the optical trap [17], so that our measurements without the trap might be less force dependent. We therefore performed intersection simulations ($n = 10$ simulations with a microtubule separation of 50nm) with different values of $\delta_{2r}$. Decreasing $\delta_{2r}$ from 1.1nm to 0.7nm eliminated termination events (Fig. S11), so we used $\delta_{2r} = 0.7$nm in our intersection simulations.

If our reasoning is correct that the decrease in $\delta_{2r}$ arises from a lack of vertical forces in our microtubule intersection experiments, then it is likely that $\delta_{2r}$ will be different in our laser trap measurements. For these, we chose an intermediate value of $\delta_{2r} = 0.9$nm. We justify a reduced value compared to the previously reported value [12] for two reasons. First, we used smaller beads in our experiments, so we would expect to have smaller vertical forces. Second, it is a challenge to simultaneously explain our observations in the laser trap that mostly 1-3 kinesin motors are engaged with a microtubule and also our observation that termination events rarely occur at intersections. An increased value of $\delta_{2r}$ and associated increase in kinesin detachment would make it easier to explain the apparent decrease in attached kinesin in the laser trap. We therefore intentionally chose a conservative value of $\delta_{2r} = 0.9$nm for our laser trap simulations.

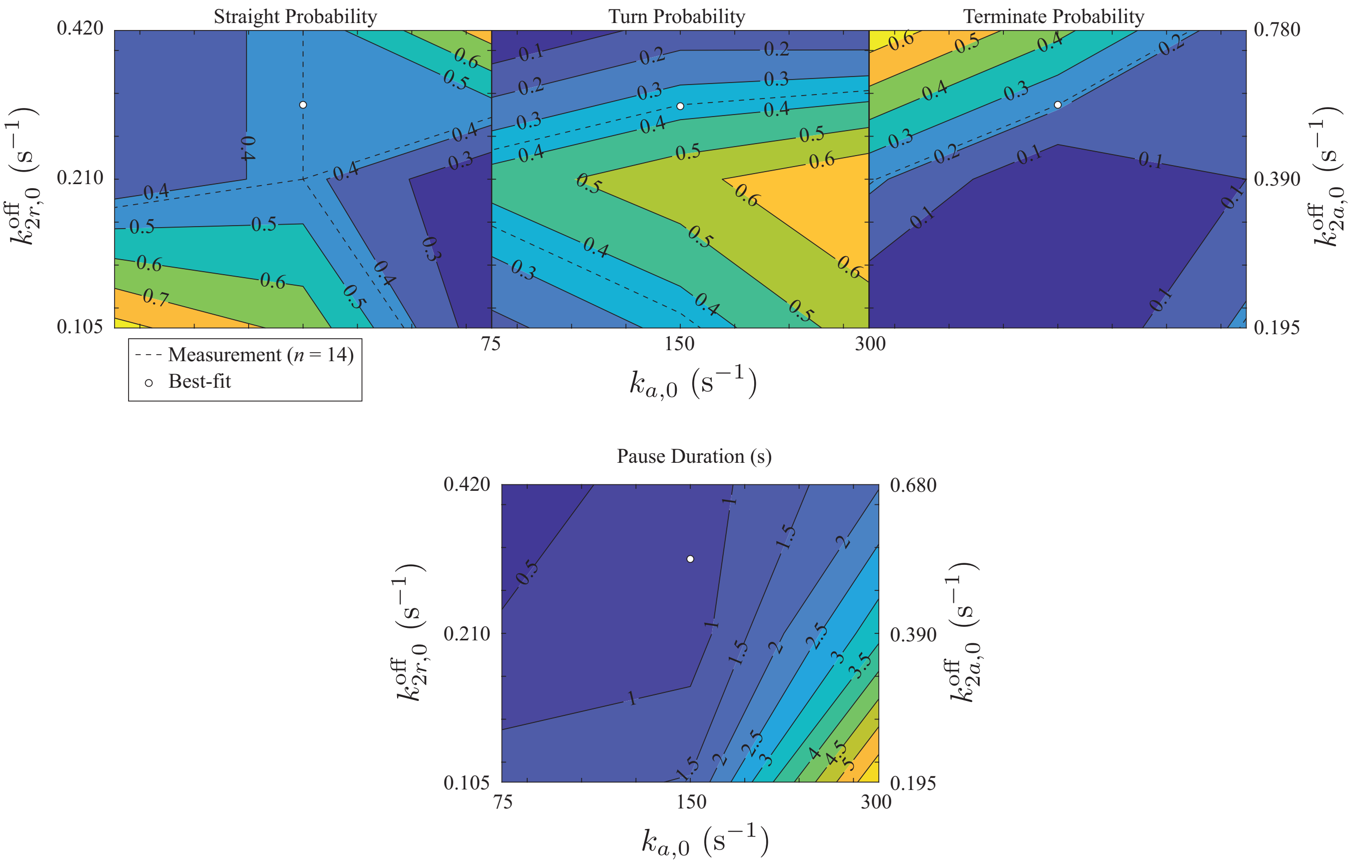


Figure S12: A sensitivity analysis of transport through microtubule intersection at a separation of $d = 100$nm suggests that model parameters are close to the best-fit parameters. Top, straight (left), turn (middle), and terminate (right) probability for different attachment (horizontal axis) and detachment (vertical axis) rates. Measured probabilities are shown as dashed lines on each plot, and the intersection of the dashed lines as a hollow dot. Bottom shows pause duration. Note that each axis is on a log scale.

#### 10.4 Sensitivity analysis

After estimating all model parameters, we then predicted our experimental measurements for the laser trap experiments and for the 3D microtubule intersection experiments. We find reasonable agreement between the model predictions and the measurements, supporting the conclusion that the model accurately captures the molecular details that occur in our experiments. However, it is unclear whether 1) we have found the optimal parameter values, 2) our results depend on our parameter estimates, and 3) our observations would change for different kinesin motors. To address these issues, we performed a sensitivity analysis.

We considered three different attachment rates, $k_{a,0} = 75, 150$, and $300\text{s}^{-1}$ and three different detachment rates $(k^{\text{off}}_{2r,0}, k^{\text{off}}_{2a,0}) = (0.105, 0.195), (0.210, 0.390)$ and $(0.420, 0.780)\text{s}^{-1}$. These values correspond to one half, one, and two times our estimates of $k_{a,0} = 150\text{s}^{-1}$ and $(k^{\text{off}}_{2r,0}, k^{\text{off}}_{2a,0}) = (0.210, 0.390)\text{s}^{-1}$. Then, for each of the 9 possible parameter combinations, we performed 10 simulations of liposomes with ten kinesin motors approaching a microtubule intersection of separation $d = 100$nm. The outcome probabilities and pause durations are shown in Fig. S12.

These simulations show that any outcome (straight, turn, terminate) can be favored by appropriately tuning the attachment and detachment rates (Fig. S12). Thus, the fact that straight outcomes were favored in our experiments and our model is not a generic outcome, but rather depends on parameter values. This is different from our previous modeling of myosin Va-based transport into actin intersections, where we always observed a bias toward going straight, regardless of parameter values [10]. This difference is likely due to a combination of 1) kinesin motors mostly proceeding along a single protofilament, as opposed to myosin Va motors which spiral around the actin filament, 2) mostly one or two kinesin motors being bound to the microtubule, as opposed to mostly three myosin Va motors being bound to actin, and 3) microtubules being larger in diameter than actin, all of which cause the crossing microtubule to present a more significant barrier to kinesin-based transport than the crossing actin filament does for myosin Va-based transport.

Looking more closely at the results, we can see that increasing or decreasing both attachment and detachment rates (i.e., moving up and to the right or down and to the left in the plots, Fig. S12) increases the probability of going straight, with a corresponding decrease in turning probability and little change in termination probability or pause duration. Alternatively, we can favor turning at the expense of going straight by increasing attachment (i.e., moving to the right in the plots, Fig. S12), which increases pause duration. Finally, we can favor termination by increasing detachment while decreasing attachment (i.e., moving up and to the left in the plots, Fig. S12), which decreases pause duration. When we plot the probabilities of going straight, turning and terminating from our simulations (obtained from all measurements with microtubule separations between 85 and 115nm, $n = 14$), we find that there is a predicted best-fit parameter combination with a similar attachment rate and a similar, but slightly ($\sim 1.4$ times) faster detachment rate than our best estimates. We therefore conclude that our parameter estimates are supported by our measurements of outcome at microtubule intersections.

#### 11 Effect of vertical force dependence

Though successful in reproducing our measurements in many respects, the model differs in a few details. One example of such a difference is intersection outcome at small separation ($d < 175$nm) and large approach angle ($\alpha > 75°$) where the measured straight:turn ratio is 1.3 while the model predicts a ratio of 11.5 (Fig. 7 of the main text). A possible explanation of this discrepancy is that, in the model, kinesin's force-dependent detachment depends only on the component of force along the microtubule and is independent of the component of force normal to the microtubule. There is, however, recent evidence that vertical forces accelerate microtubule detachment for kinesin-1 [17]. This effect was discussed previously (Section 10.3), to motivate a smaller dependence on forces directed along the microtubule in the unbinding rate of kinesin than measured at the single molecule level [12]. However, by modeling this effect as an overall decrease in force dependence, we neglect the fact that, in the intersection simulations, this effect may differ for different approach angles.

When liposomes approach an intersection at a large approach angle, the kinesin molecules are likely attached to the side of the starting microtubule (Fig. S13A, top left). If a tug of war occurs, the kinesin molecule(s) on the crossing microtubule apply a force oriented vertically from the point of view of the kinesin molecule(s) on the starting microtubule (Fig. S13A, top right). Conversely, when liposomes approach an intersection at a small approach angle, the kinesin molecules are likely attached to the top of the starting microtubule (Fig. S13A, bottom left). The net force of the kinesin on the crossing microtubule would be oriented more laterally (i.e., to the left or right) from the perspective of kinesin on the starting microtubule (Fig. S13A, bottom right).

It therefore seems an intriguing hypothesis that some of the differences between model and experiment might be lessened or removed if we include vertical force dependence in our model. Unfortunately however, the precise vertical force dependence of detachment in kinesin has not been measured, although it is clear that larger vertical loads promote detachment more rapidly than smaller ones [17]. So, while we could not test this hypothesis by implementing vertical force-dependence in our model using previous measurements, we could test the hypothesis by adding vertical force-dependence to our model. The hypothesis then

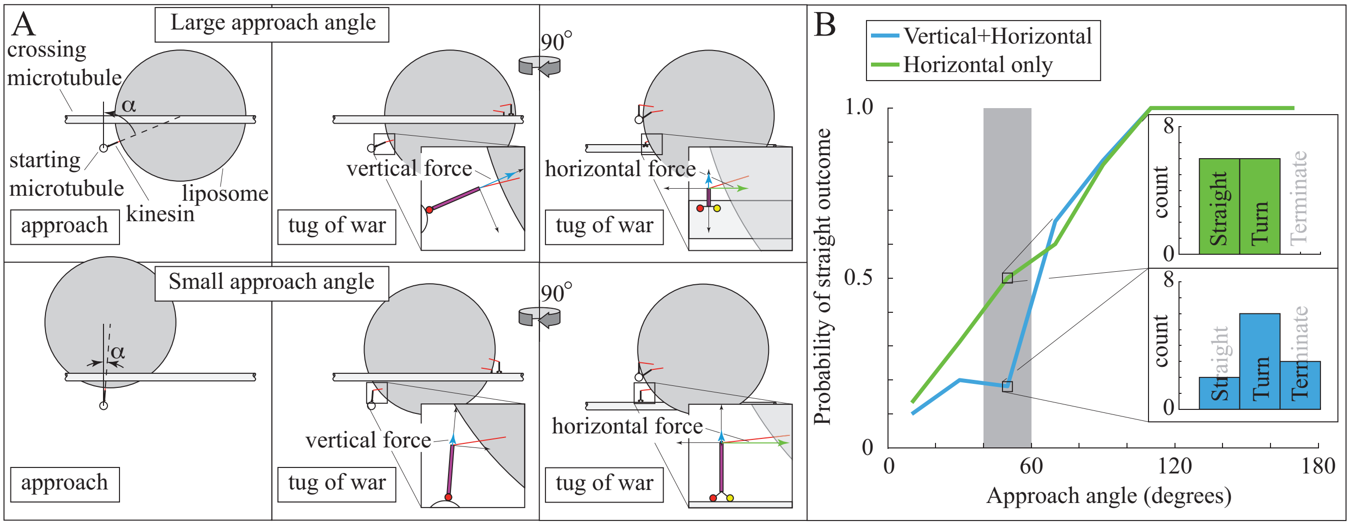


Figure S13: Vertical force dependence of detachment decreases the probability of proceeding straight through a microtubule intersection at an intermediate approach angle. A. Cartoons of liposomes approaching a microtubule intersection at a large approach angle and subsequent large vertical force during a tug of war (top) and a small approach angle and subsequent small vertical force during a tug of war (bottom). Note that the middle panel shows the full vertical force component (blue), while the horizontal component is absent because it points out of the page. The right panel shows the full horizontal force component (green), while the vertical component appears shorter, particularly in the top panel, because it points partly out of the page. B. The probability of going straight through a microtubule intersection for simulations with (blue) and without (green) vertical force dependence, for microtubule separations of $d = 50, 75$ and 100nm. The simulations predict similar probabilities, except for intermediate approach angles ($40° < \alpha \leq 60°$, shaded). A more detailed look at the outcomes for these approach angles (inset) shows that turning probability is unaffected by the vertical force dependence, decreasing the straight:turn ratio.

predicts we should see an increase in turning probability at larger approach angles.

To implement vertical force-dependence in our model, we modified detachment from state 2 in the model (Fig. S9C):

$$k_2^{\text{off}} = \begin{cases} k_{2r,0}^{\text{off}} e^{(\delta_{2r}|F|+\delta_n|F_n|)/k_BT} & : \text{ resistive force} \\ k_{2a,0}^{\text{off}} e^{\delta_{2a}|F|/k_BT} & : \text{ assistive force} \end{cases}$$

Where we have modified the force dependence for resistive force to include a force dependence $\delta_n$ that increases detachment for the component of force normal to the surface of the microtubule $F_n$. We performed simulations using the same value of $\delta_{2r} = 0.7$nm as in our intersection simulations and $\delta_n = 0.5$nm. We performed these simulations identically to the intersection simulations (Section 9), but only considered small microtubule separations, $d = 50, 75$, and 100nm. Interestingly, we find that simulations with vertical force dependence are similar to those with only horizontal force dependence for small and very large approach angles (Fig. S13). However, for approach angles between $40^\circ < \alpha \leq 60^\circ$, the simulations with vertical force dependence predict a much smaller probability of going straight (less than 20%) compared to the simulations without (50%, Fig. S13). In fact, the straight:turn ratio decreases from 1 to 0.33 in the presence of vertical force dependence (Fig. S13, inset). Consistent with our hypothesis, vertical load dependence increases the turning probability of liposomes approaching a microtubule intersection at larger approach angles.

This result suggests that a model with vertical load dependence could agree with our measurements better than our current model. However, testing this idea by fitting the model to our measurements would require redoing the analysis presented in Section 10 of this supplement for a range of different normal force dependencies, $\delta_n$. The computational cost of such an approach is too large and, in any case, the predicted value of $\delta_n$ would need to be experimentally verified. It will therefore be critical for the field to independently define the vertical and horizontal load sensitivity of kinesin motors so that we may understand their behavior in a 3D environment more clearly.
